# Arkadia via SNON enables NODAL-SMAD2/3 signaling effectors to transcribe different genes depending on their levels

**DOI:** 10.1101/487371

**Authors:** Jonathon M. Carthy, Marilia Ioannou, Vasso Episkopou

## Abstract

How cells assess levels of signaling and select to transcribe different target genes depending on the levels of activated effectors remains elusive. High NODAL-signalling levels specify anterior/head, lower specify posterior, and complete loss abolishes anterior-posterior patterning in the mammalian embryo. Here we show that cells assess NODAL-activated SMAD2 and SMAD3 (SMAD2/3) effector-levels by complex formation and pairing each effector with the co-repressor SNON, which is present in the cell before signaling. These complexes enable the E3-ubiquitin ligase Arkadia (RNF111) to degrade SNON. High SMAD2/3 levels can saturate and remove SNON, leading to derepression and activation of a subset of targets (high targets) that are highly susceptible to SNON repression. However, low SMAD2/3 levels can only reduce SNON preventing derepression/activation of high targets. Arkadia degrades SNON transiently only upon signaling exposure, leading to dynamic signaling-responses, which most likely initiate level-specific cell-fate decisions. Arkadia-null mouse embryos and Embryonic Stem Cells (ESC) cannot develop anterior tissues and head. However, SnoN/Arkadia, double-null embryos and ESCs are rescued confirming that Arkadia removes SNON, to achieve level-dependent cell-fates

**One Sentence Summary:** Signaling intensity induces equivalent degradation of a transcriptional repressor leading to level-dependent responses.

## Introduction

NODAL is a morphogen and a member of the transforming-growth-factor-beta (TGF-β)-family ligand(*1*). It is transduced via C-terminal phosphorylation of SMAD2 and SMAD3 class of transcriptional effectors (SMAD2/3; hereafter refers to signaling-activated forms). SMAD2/3 complex with SMAD4 and translocate to the nucleus where they bind to DNA to initiate transcription(*2–5*). SMAD2/3 target-gene specificity is regulated by interactions with other transcription partner-factors present in the cell(*4, 6*). However, genetic experiments in mouse(*7*) and gain of function experiments in *Xenopus* and *zebrafish(1, 8)* show that during gastrulation different NODAL-signaling levels (intensities) activate distinct set of targets to establish various cell-fates along the anterior-posterior axis of the embryo. However, mechanisms that allow cells to assess NODAL-signaling levels and activate level-specific transcriptional responses remained unknown(*9*).

Gastrulation in mice is initiated by delamination of the epiblast epithelium at the prospective posterior end, followed by elongation of the delamination front towards the center of the embryo leading to Primitive Streak (PS) formation(*10*). Cells express *Nodal* only when they ingress within the PS, and turn it off when they migrate out laterally specified into precursors of specific tissues (germ layers)(*7*). In addition, these migrating precursors possess information about the prospective position that they will occupy along the anterior posterior axis of the embryo. Gradual reduction of the SMAD2/3 effector-alleles using genetics in mouse causes loss of cell-fates starting with those that derive from the elongating front of the PS (anterior-PS). Embryos with more advanced loss of effector-allele lose additional cell fates that derive from more posterior PS such as somites(*11*). These experiments suggest that graded NODAL-signaling is responsible for patterning the PS precursors with anterior-PS depending on maximum levels. Several NODAL targets operating during development have been identified however, it remained unclear whether they are activated by high or low NODAL signaling.

We previously showed that the RING-domain E3 ubiquitin ligase, RNF111 (Arkadia), is essential for the formation of anterior-/head tissues, but not of posterior, by a mechanism that depends on NODAL(*12–14*). Complementary experiments in *Xenopus* animal-cap assays showed that anterior cell fates are induced by co-injection of *Nodal* and *Arkadia* mRNAs but not by the same amount of *Nodal* or *Arkadia* injected alone, confirming that Arkadia potentiates NODAL signaling(*13*). Biochemical and cell culture experiments showed that Arkadia enhances signaling by targeting for degradation (a) the inhibitory SMAD7(*15*), which attenuates signaling mainly at the receptor level(*16, 17*), and (b) the nuclear co-repressors SNON and SKI(*18, 19*), which bind to SMAD2, SMAD3 or SMAD4 and suppress their transcriptional responses(*20*). Notably, Arkadia has been shown to degrade SNON only when it is bound to SMAD2 or SMAD3(*18*). Whether the above substrates mediate the function of Arkadia in the embryo had not been determined.

Here we show that *Ark*−/− embryos, which lack anterior/head structures, are rescued by genetic removal of *SnoN*, but not of *Smad7*, while embryos with single deletions of either *SnoN* (*SnoN−/−*) or *Smad7* (*Smad7*−/−) are normal. Similarly, *Ark*−/− ESCs cannot differentiate towards anterior cell-fates and exhibit defective activation of a subset of SMAD2/3-targets, but these are rescued in double-null *Arkadia* and *SnoN* ([*Ark/SnoN*]−/−) ESCs. We also show that Arkadia degrades the SNON-SMAD2 complex to achieves derepression within the first hour of signaling, and that SNON reduction is transient and proportionate to SMAD2/3 levels and derepression. Collectively, the above data reveal how cells assess NODAL-signaling levels and convert each of them to specific gene-expression patterns and cell fates.

## Results

### Arkadia removes SNON to induce anterior in ESCs and embryos

Cell-fates derived from the anterior-PS include the anterior definitive endoderm (ADE); the node, which is a structure forming at the end of the fully elongated PS after ADE migration; and the node-derived mesendoderm (ME), which is a cord-like structure extending along the midline joining the node with the ADE(*7*). ESCs differentiate towards ADE-ME with Activin-A (Activin; NODAL-like ligand) treatment(*21*). To address whether SNON is the substrate that mediates Arkadia’s function we used *WT* and *Ark*−/− ESC lines derived from mouse embryos that carry the *Hex-Gfp* transgene, an ADE-ME reporter(*22*). *WT* ESC-lines (n=3) differentiate normally towards ADE-ME with Activin, while none of the *Ark*−/− lines (n=3) do so (Fig. **1A**). However, deletion of *SnoN* via CRISPR/Cas9 in *Ark−/−* ESCs [*SnoN/Ark*]−/−, (n=10 clones; Fig **S1A**), rescues their ability to differentiate towards ADE-ME (Fig 1a). We confirmed the ADE-ME differentiation by quantitative PCR (qPCR) using the definitive endoderm marker *Cxcr4*(*23*), the ME marker *Foxa2*(*24*)(Fig 1B), and by western blot with GFP antibody (Fig. **S1B**). *SnoN*−/− ESC lines (n=3) differentiate normally towards ADE-ME (Fig. **S1C** and **S1D**), excluding the possibility that loss of SNON leads to excessive ADE-ME differentiation and it is responsible for the rescue observed in [*SnoN/Ark*]−/− cells.

**Fig. 1:**
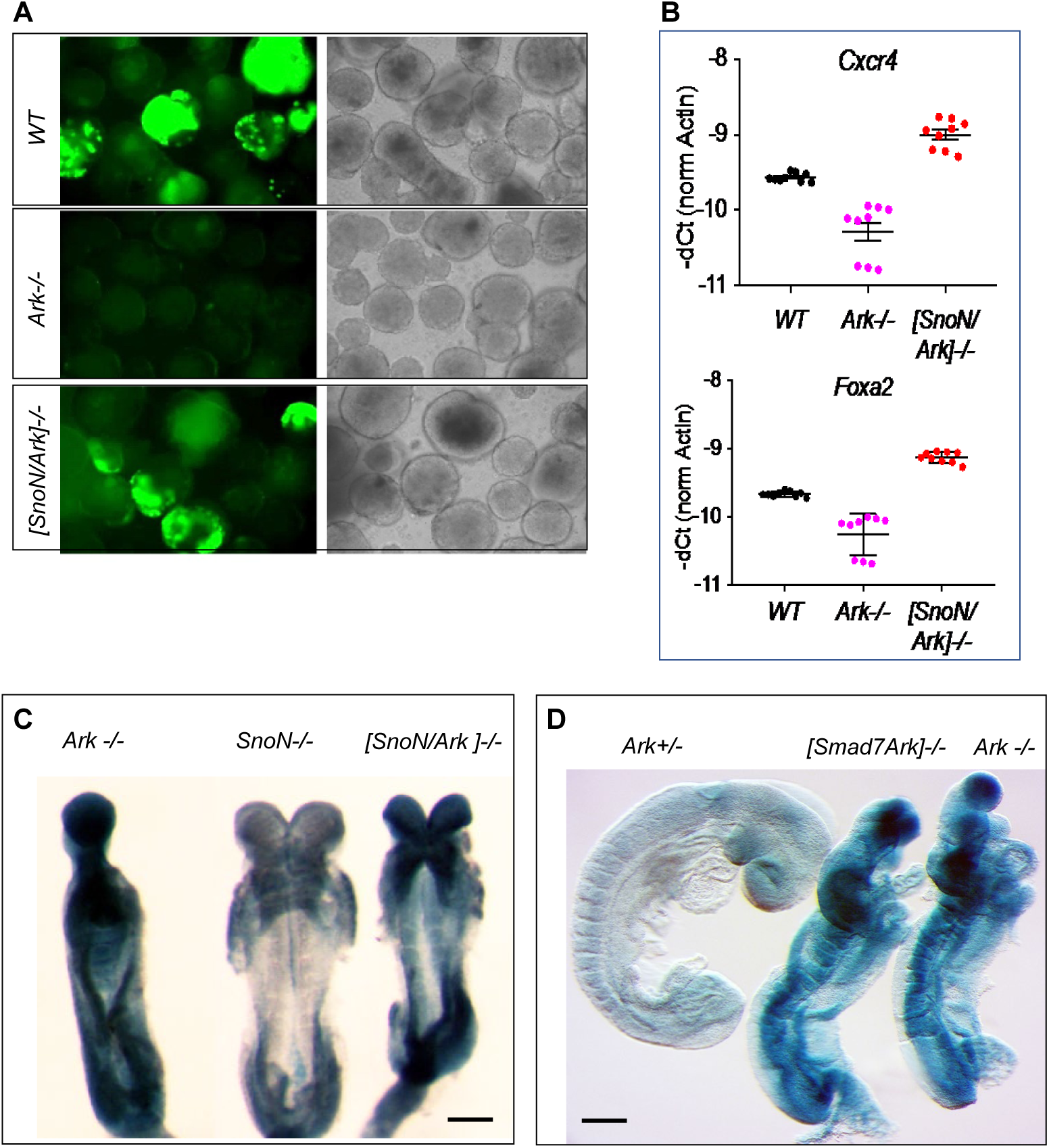
Arkadia removes SNON in ESCs and embryos. (**A**) *Hex-Gfp* ESCs differentiated towards ADE. Fluorescence (left) and bright field (right) images. (**B**) QPCR expression analysis of ADE/ME markers. (**C**) X-gal stained 6-8-somite E8.5 embryo from [*SnoN−/−;Ark+*/-]x[*SnoN−/−;A*rk+/-] cross. Out of 63 embryos E8.5-9 analyzed, 18 were [*SnoN/Ark*]−/−, of which 16 seemed fully and 2 partially rescued (88% rescue). The rest of the embryos were of the parental genotype and were normal. (**D**) X-gal stained E9.5 embryos (>10 somites) from [*Smad7+/-;Ark+*/-]x[*Smad7+/-;A*rk+/-] cross. Out of 207 embryos analyzed, 48 (23%) exhibited *Arkadia*-null phenotype of which 15 (7.3%) were [*Smad7/Ark*]−/− (fewer than expected due to early resorptions). The rest of the embryos were single or double heterozygous and looked normal. *SnoN*-null and *Ark*-null alleles express *β-galactosidase* via their endogenous(*12*). Genotypes are as indicated; scale bar is 200μm.

In the embryo the brain/head is induced by ADE and the adjacent ME domain (prechordal plate)(*25*), which migrate from the anterior-PS across and opposite, where they induce anterior neuroectoderm (brain) on the overlaying ectoderm layer(*25*). To address whether removal of *SnoN* from *Ark*−/− embryos can rescue their phenotype, we used mice, homozygous for *Skil*^*tm2Spw*^(*26*), which are viable and fertile. We confirmed that these embryos are indeed *SnoN-* null (*SnoN−/−*) using whole-mount *in-situ* hybridization (WISH) with *SnoN* probe (Fig. **S2A-C**). The absence of an embryonic phenotype in *SnoN−/−* mice is puzzling however, we found that SKI, the close homologue of SNON, is upregulated in *SnoN*−/− embryos (Fig. **S2E**) suggesting compensation. Furthermore, using WISH with the ADE marker(*27*) *Cerberus (Cer*) we did not find any obvious difference in size or timing of ADE formation in *SnoN*−/− embryos (Fig. **S2D**). SNON is one of several feedback mechanisms that regulate fluctuations of NODAL-signaling intensity. Together these mechanisms increase the robustness of signaling regulation and most likely prevent dysregulation and phenotypes in the absence of SNON *in vivo*.

Consistent with the rescue that we observed in ESCs differentiation, [*SnoN/Ark*]*−/−* embryos also exhibit rescue of ADE, node-ME development along with midline and brain/head formation (Fig. **1C**) with high penetrance (>88%). Arkadia also degrades the NODAL-signaling inhibitor SMAD7(*15*), which mainly mediates the degradation of the TGFβ-signaling receptors(*16, 17*) and therefore functions differently than SNON and SKI. *Smad7*−/− (*28*) mice, do not exhibit patterning defects during gastrulation and embryos develop normally, with some being born and surviving to adulthood. We generated [*Smad7/Ark*]−/− embryos and found that none was rescued as they are indistinguishable from the phenotype of *Ark−/−* embryos (Fig. **1D**). Therefore, SNON, and not SMAD7, is the major substrate that mediates Arkadia’s function during gastrulation.

We examined the extend of the rescue in *[SnoN/Ark]−/−* embryos using WISH with various makers: *Cer* probe showed that *[SnoN/Ark]−/−* embryos form ADE comparable to the *WT* and *SnoN*−/− embryos while *Ark*−/− do not (Fig. **2A**). *Six3* (*29*) showed that *[SnoN/Ark]−/−*, *WT* and *SnoN*−/− embryos develop forebrain while *Ark*−/− do not (Fig. **2B**). Both *FoxA2*(*24*) (Fig. **2C**) and *Shh*(*30*) markers showed that unlike Ark−/−, *[SnoN/Ark]−/−, WT* and *SnoN*−/− embryos, form full length midline all the way to the brain (Fig 2d). The above embryo analysis confirms rescue of anterior-PS tissue development in *[SnoN/Ark]−/−* embryos. *Shh* marks also the hindgut endoderm(*11*) revealing that it forms in *Ark*−/− embryos (arrows in Fig. **2D**) most likely because it derives from posterior- and not anterior-PS. However, *[SnoN/Ark]−/−* pups (n>50 litters) are not born indicating that later development cannot proceed in the absence of both Arkadia and SNON. We cannot exclude the possibility that SKI also mediates the function of Arkadia but to lesser extent. The most likely explanation for the above results is that a subset of NODAL-SMAD2/3 target genes responsible for the anterior-PS cell-fates are suppressed by SNON and depend on Arkadia and signaling for derepression and subsequent activation.

**Fig. 2:**
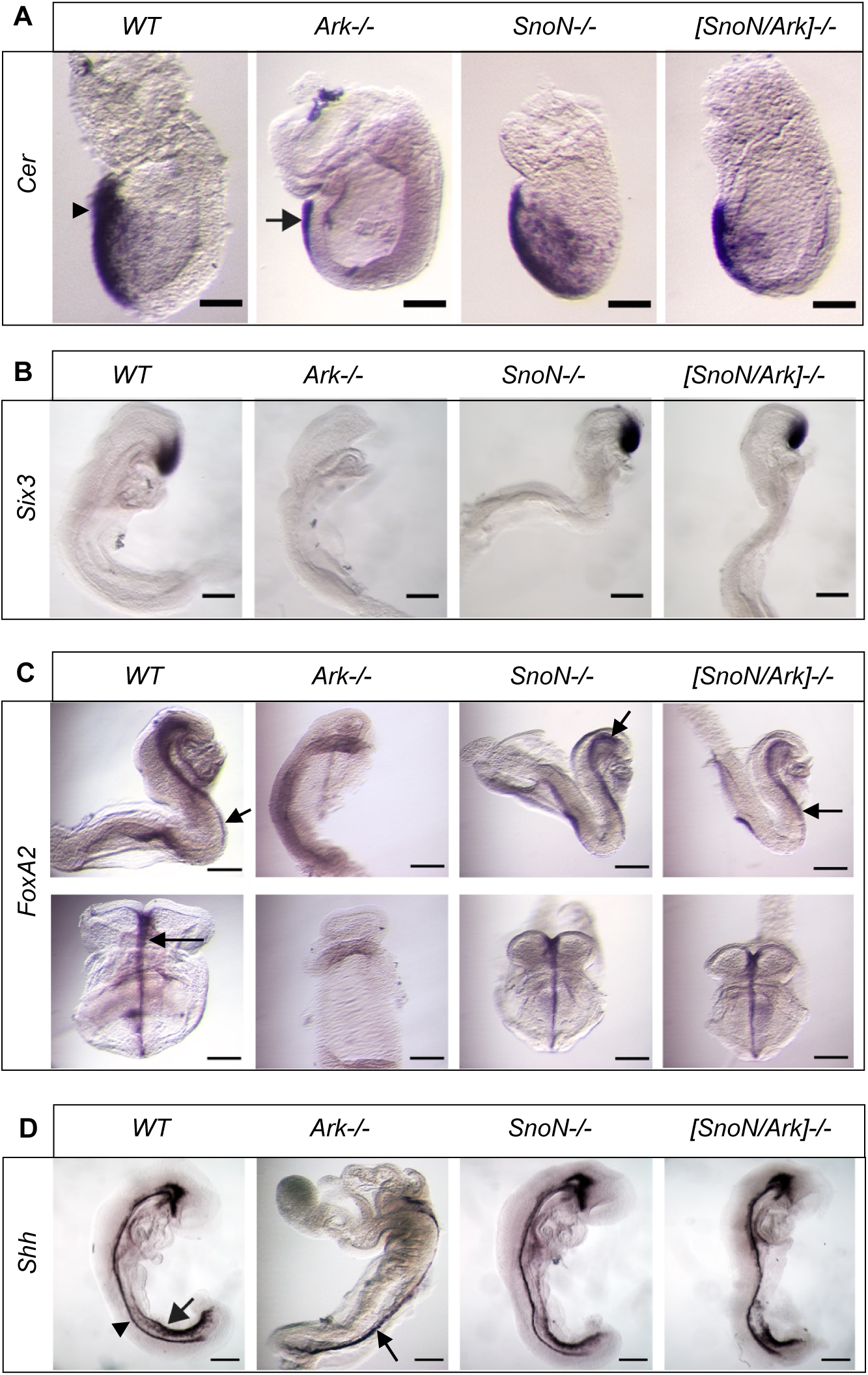
Arkadia functions via SNON in the embryo. WISH with markers as indicated: (**A**) E7.5 embryos stained with *Cer* are shown with anterior on the left and scale bar 100µm. *Ark*−/− embryos are known to lack ADE (arrowhead) but retain the extraembryonic anterior visceral endoderm (AVE) domain (arrow)(*14*). (**B**) Lateral view of E8.5 embryos stained with *Six3* is shown. *Six3* expression is rescued in [*SnoN/Ark*]−/− embryos. (**C**) *Foxa2* stained E8.5 embryos with lateral (top) and ventral/frontal (bottom) views are shown with arrows pointing the ME. (**D**) Lateral view of *Shh* stained E9.5 embryos are shown with arrows pointing the hindgut and arrowheads the ME. Scale bar: 100µm (**A**); 200µm (**B-D**).

### Arkadia derepresses SNON-repressed genes

To identify Arkadia-dependent NODAL target genes we used two different *WT* and two *Ark−/−* ESC-lines (undifferentiated) and performed duplicate time course experiments. We shut-off endogenous signaling with the TGFβ-receptor kinase inhibitor (SB431542; SB) and then harvested the ESCs for protein and transcriptomic analysis either without any other treatment (T0) or after subsequent treatment with Activin for 3-(T3) or 6-hours (T6). Western blot revealed that SNON and SKI repressors are present at high levels prior to signaling and are degraded upon signaling initiation by Arkadia (Fig. **3A**). The RNA-sequencing data have been deposited in NCBI Gene expression Omnibus (accession number: GSE118005).

**Fig. 3:**
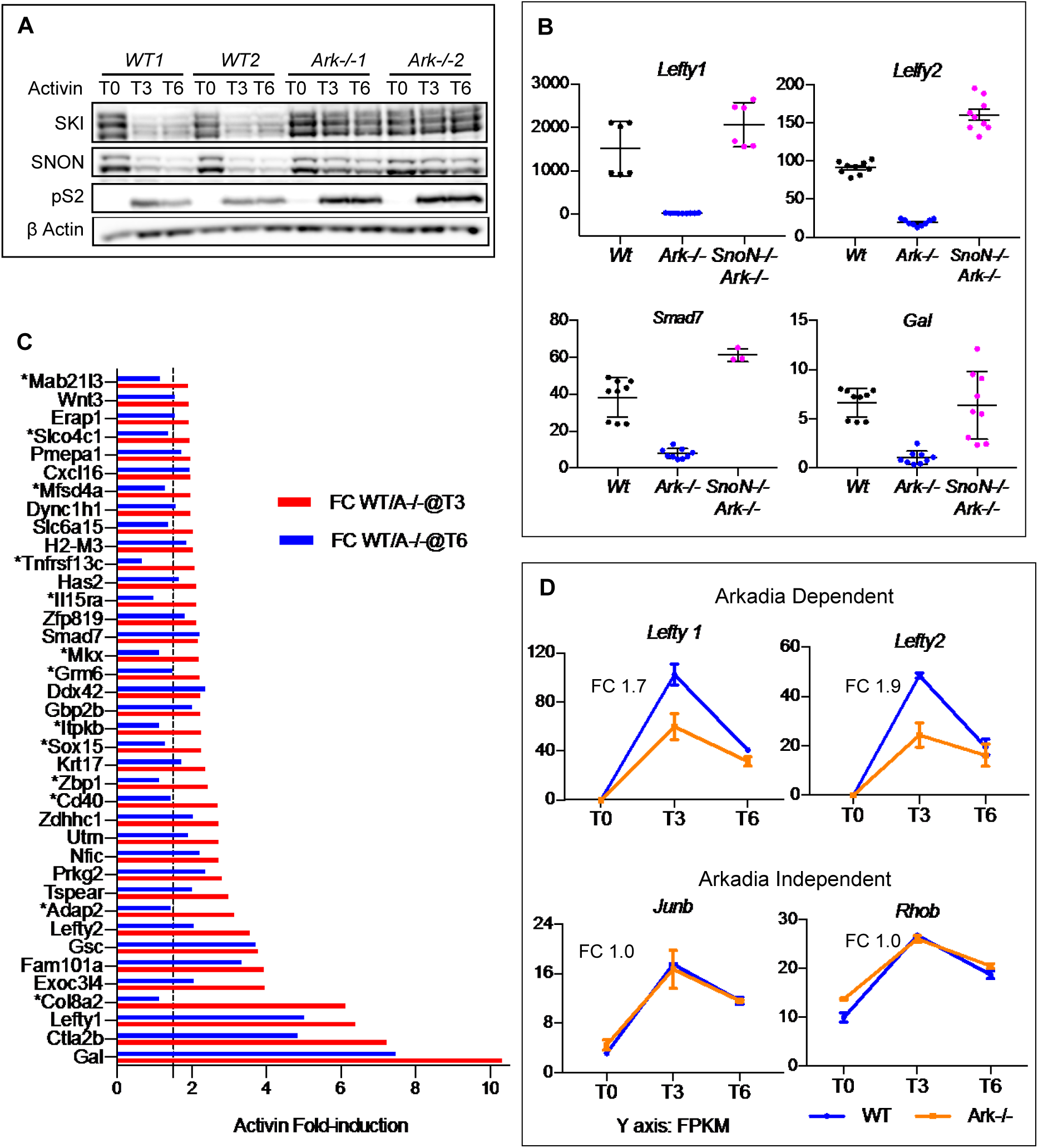
Arkadia derepresses SNON-repressed genes. (**A**) Western blot analysis of ESCs treated prior to harvesting for RNA-sequencing. Treatment, time and antibodies are as indicated (pS2: anti-SMAD2). (**B**) QPCR analysis of representative Arkadia-dependent targets in ESCs with genotype as indicated, pre-treated with SB and then with Activin for 3-hours. (**C**) A graph comparing Activin induction difference (FC) between *WT* and *Ark*−/− ESCs at T3 and T6 of 38 Arkadia-dependent genes (all with ≥1.9-fold activin induction difference between *WT* and *Ark-*/-ESCs at T3). Asterisks indicate genes showing reduced difference at T6 below ≤1.5-fold difference (dotted line). (**D**) Expression graphs of representative Arkadia-dependent and - independent SMAD2/3-targets from the RNA-sequencing results. The low activation of Arkadia-dependent targets at T3 in Ark−/− ESCs is probably due to the mild SNON/SKI reduction observed in (**A**) by other ubiquitin ligases known to degrade them(*20*) causing derepression.

We found 262 genes with ≥2-fold Activin induction in both *WT* ESC-lines at T3 (Table **S**) including several known SMAD2/3 targets(*31*) are included in this list, providing confidence that they are Activin-induced. 52 of these genes exhibit ≥1.7-fold downregulation in *Ark−/−* ESCs compered to *WT* at T3, and were considered as Arkadia-dependent (red font, Table **S**). We then examined the expression of some of these genes in [*SnoN/Ark]−/−* ESC-lines generated by CRISPR/Cas9, and found that their expression is restored at T3 (Fig. **3B**). These genes exhibit *WT* levels of expression in *SnoN*−/− ESCs (Fig. **S3A**) excluding the possibility that there is an over-induction/expression of target genes in the absence of SNON that causes the rescue in [*SnoN/Ark*]−/− ESCs. This confirms that SNON represses a subset of Activin-induced target genes and that Arkadia removes SNON to allow their expression.

SNON was thought to inhibit broadly SMAD2/3-transcriptional responses because it is recruited to targets by binding to SMAD2, SMAD3 or SMAD4(*20*). However, the above data show that SNON does not suppress equally all SMAD2/3 target genes. Although it is unknown what determines the susceptibility of target to SNON repression, the data show that highly susceptible to SNON repression targets (therefore highly dependent on Arkadia) tend to be silent in the absence of signaling (Fig. **S3B**). This suggests that the susceptibility to SNON repression is influenced by chromatin and expression potential of targets prior to signaling activation. The finding that SNON is present at high levels in the absence of signaling (Fig. **3A**) together with the fact that SMAD4 can bind to SNON and also to the same DNA element as SMAD2/3, suggest that SMAD4-SNON complexes silence SMAD2/3-targets prior to signaling. Furthermore, unlike SMAD2/3, SMAD4 does not have a transactivation domain and cannot recruit co-activators and histone acetylases to its targets, suggesting that SMAD4-SNON are more likely to mediate target gene repression. Recent findings support the above reasoning by showing that SMAD4-SKI represses the master Th17 differentiation gene, RORγt, which depends on TGFβ signaling for derepression(*32*).

Notably, Arkadia-dependent, SMAD2/3 targets are downregulated by≥1.7-fold in *Ark*−/− ESCs compared to *WT* at T3, while at T6 this deficiency is reduced (Fig. **3C** and **3D**; and Fig. **S4A**). This is caused by a significant downregulation of several Activin-induced targets in *WT* ESCs at T6 compared to T3 irrespective of whether they are Arkadia dependent or not, reflecting negative feedback and signaling dynamics. Collectively, the above experiments identified a subset of Activin/NODAL-induced targets susceptible to SNON repression and showed that derepression/activation of these targets by Arkadia is transient and occurs within the first 3 hours of ligand stimulation.

### Arkadia degrades SMAD2/3-SNON complexes

Arkadia is known to function by degrading the repressors SNON and SKI in the presence of TGFβ signaling(*19*) and that the interaction of SNON with SMAD2 or SMAD3 is required for SNON degradation(*18*). Consistent with this in *WT* and *Ark*−/− ESCs SNON and SKI are high in the absence of signaling and are degraded within the first hour upon TGFβ-ligand treatment only in *WT* ESCs but not in *Ark*−/− (Fig. **4A**). Similar to other cell lines(*18*), SNON levels, unlike those of SKI, recover over time with continuous signaling in *WT* ESCs. This coincides with the downregulation of the Activin induced transcriptional responses observed in our RNA-sequencing experiments at T6 compared to the earlier time of T3 (Fig. **3C** and **3D**; and Fig. **S4A**). Unlike *Ski, SnoN* gene is upregulated directly by SMAD2/3 (>3.2-fold; Table **S**)(*33, 34*), and this possibly overwhelms Arkadia, which is known to be unstable(*14*) and therefore contributes to the SNON resistance.

**Fig. 4:**
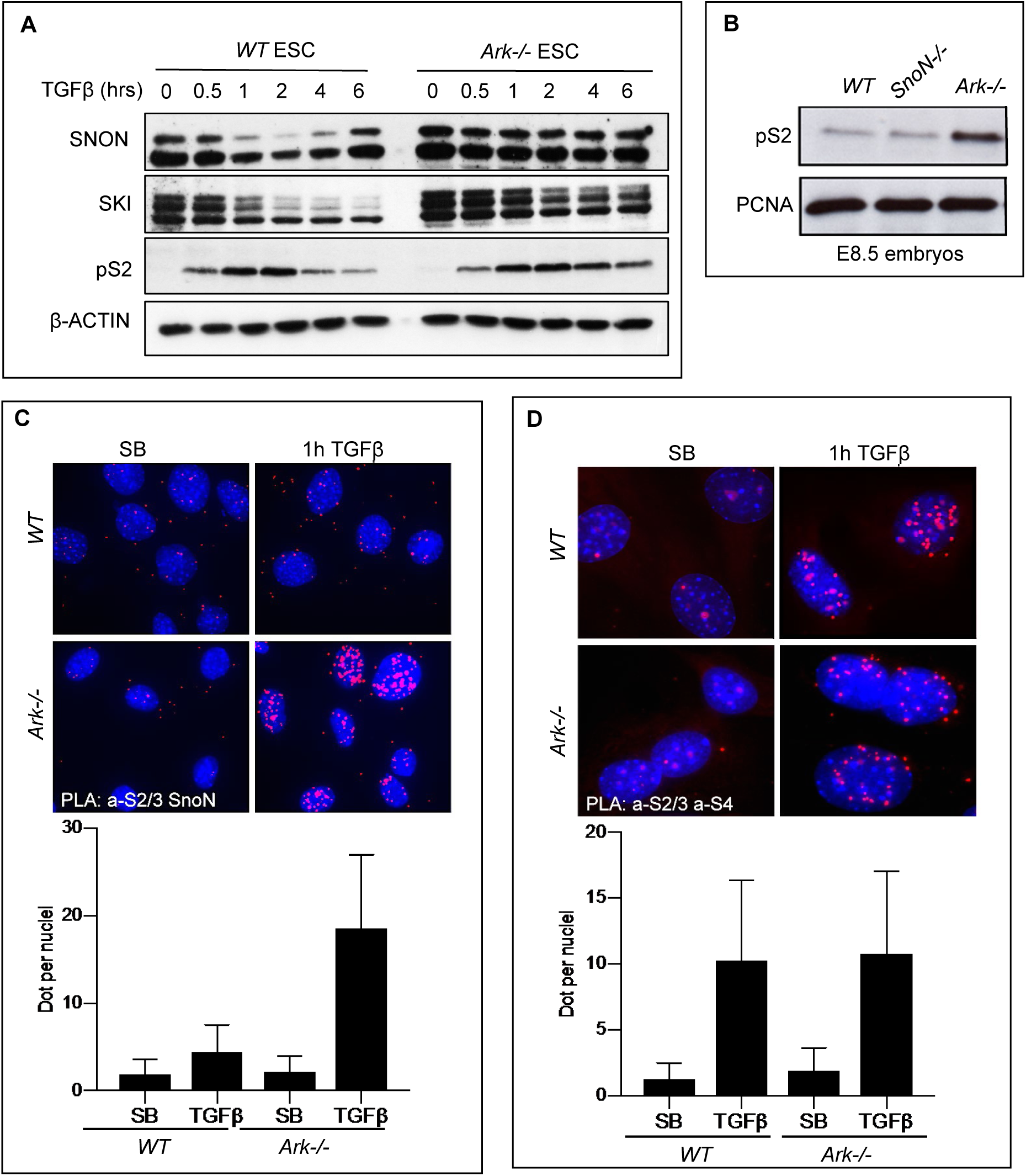
SMAD2/3 via Arkadia drive SNON degradation. Western blot analysis with antibodies as indicated, on extracts from (**A**) ESCs treated with TGFβ ligand over time or SB (“0”) and (**B**) from embryos with genotypes as shown. The full recovery of SNON levels observed here at 6hrs is not seen in Fig. **3A** most likely due to different culturing conditions (see methods). (**C** and **D**) Images of PLA experiments (top) and quantitative analysis (bottom) with antibodies, total-SMAD2/3 (S2/3) and SNON in (**C**), and S2/3 and SMAD4 (S4) in (**D**). Treatment and genotypes of MEFs are as indicated.

Furthermore, the above experiments showed that in the absence of Arkadia SMAD2, like SNON, accumulates in ESCs (Fig. **3A**, Fig. **4A**) and embryos (Fig. **4B**) possibly in a complex suggesting that Arkadia degrades both. This is supported by our previous study showing that in ESCs Arkadia enhances signaling by interacting directly, ubiquitinating and degrading SMAD2 and SMAD3(*14*). However, at that time, it was thought that Arkadia degrades the transcriptionally active fraction of SMAD2 which is bound to SMAD4. To address this further, we performed Proximity Ligation Assays (PLA)(*35*) to visualize SMAD2-SNON (Fig. **4C**) and SMAD2-SMAD4 (Fig. **4D**) complexes in mouse embryonic fibroblasts (MEFs) treated with either SB or with ligand. The data show high accumulation of SMAD2-SNON complexes only in the absence of Arkadia, and equal SMAD2-SMAD4 levels in *WT* and *Ark*−/− cells supporting that Arkadia degrades SMAD2-SNON and not SMAD2-SMAD4 complexes. Furthermore, in *WT* cells ligand treatment for one hour seems to be sufficient to eliminates all SMAD2-SNON complexes (Fig. **4C**). This suggests that the surge of SMAD2 generated upon signaling initiation pairs with SNON enabling Arkadia to degrade this complex, leading to derepression/activation of the SNON repressed targets. However, because SNON becomes resistant to Arkadia over time with signaling degradation and derepression are transient.

### Signaling intensity drives proportionate SNON reduction and level-dependent responses

The above pairing of SMAD2-SNON and their degradation by Arkadia seems to perform a comparison of the incoming SMAD2 protein levels relatively to those of SNON which are present in the cell prior to signaling. Therefore, this mechanism most likely provides an assessment of ligand-level exposure and signaling activation for each cell. To investigate whether in the presence of Arkadia the levels of signaling drive proportional SNON-depletion, we assessed SMAD2 and SNON levels downstream specific signaling levels in *WT* ESCs. To bypass the endogenous expression of various TGFβ ligands in ESCs, we controlled signaling activation at the receptor level using SB receptor inhibitor. For this we stimulated cells for one hour using fixed amount of Activin (Fig. **5A**) or TGFβ (Fig. **S4C**) ligands but with different concentration of SB to achieve different levels of signaling-activation. Quantification of the blot showed inverse correlation between SMAD2 and SNON levels with SMAD2:SNON ratio (Fig. **5B**) directly proportionate to the levels of signaling. These experiments indicate that in *WT* ESCs, different signaling levels achieve an equivalent SNON reduction and confirms that SNON is the cellular standard that signaling levels are evaluated on.

**Fig. 5:**
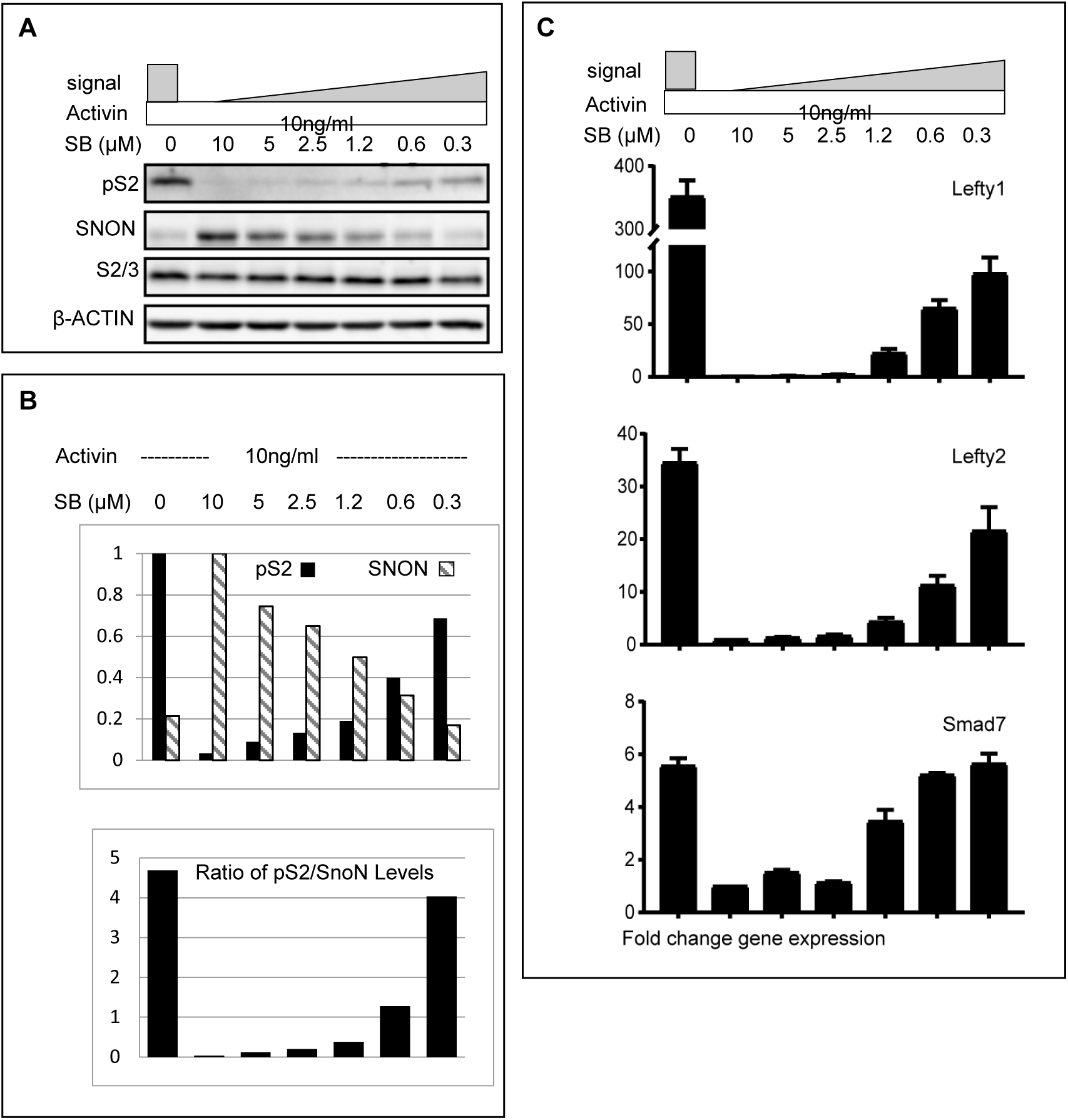
SMAD2/3 levels drive derepression. (**A**) Western blot analysis on *WT*-ESCs treated as indicated for an hour. (**B**) Quantification of pS2 and SNON proteins (top), and the pS2/SNON ratio (bottom) from the above blot. (**C**) QPCRs expression analysis for known SMAD2 target genes in *WT* ESCs treated as indicated for two hours.

To address whether the SMAD2:SNON-protein ratio correlates with the derepression and expression of known SMAD2/3 targets, we used the same Activin/SB treatment and performed qPCR analysis of representative NODAL targets. We found that the NODAL antagonists *Lefty1* (*36*) and *Lefty2* (*37*), are not activated under low levels of signaling, and they require maximum signaling for their peak activation (Fig. **5C**). Interestingly, they are not expressed at all in the absence of signaling. On the contrary, *Smad7* exhibits a low/basal level of expression in the absence of signaling and is upregulated to reach peak expression under moderate levels signaling (Fig. **5C**). These results show that each gene exhibits peak response at different SMAD2:SNON-protein ratios (compare Fig. **5B** and **5C**) suggesting variable susceptibility to SNON repression. Therefore, *Left1* is more susceptible to SNON than *Lefty2*, while *Smad7* is mildly suppressed. In the embryo, *Lefty1* is not normally expressed in the PS but *Lefty2* is. WISH showed reduced expression of *Lefty2* in *Ark*−/− embryos (Fig. **S5**) possibly due to partial derepressed by other ubiquitin ligases such as Smurf2 and APC known to also degrade SNON/SKI(*20*). This raises the possibility that a subset of SNON/SKI repressed targets is derepressed by these ubiquitin ligases in the absence of Arkadia.

Therefore, SMAD2 levels control SNON-protein reduction via Arkadia leading to derepression and expression of the SNON-repressed NODAL targets in a level-dependent fashion.

Furthermore, the variable susceptibility to repression of the NODAL targets most likely leads to unique transcriptional responses downstream of graded signaling.

## Discussion

Here we present several lines of evidence including in ESCs and embryos supporting that high NODAL-signaling targets are susceptible to SNON repression and depend on Arkadia and high signaling for SNON removal and derepression. We identified Arkadia-dependent NODAL-targets in ESCs and showed that they exhibit variable susceptibility to SNON repression and that their activation occurs under distinct levels of SNON reduction downstream equivalent SMAD2 levels. Therefore, graded NODAL signaling causes variable SNON-reduction and derepression leading to distinct expression patterns.

We showed that SNON becomes resistant to Arkadia over time under continuous NODAL-ligand exposure and that this reinstates target-gene repression in ESCs, suggesting that the initiation of high NODAL-signaling cell-fates, such as ADE, can be reversed with signaling duration. The fact that ADE cells migrate fast out and opposite of the PS and at the same time activate stable expression of *Cerberus*, a triple antagonist of NODAL, BMP and WNT(*38*), suggests that the consolidation of ADE fate requires transient exposure to high signaling. These findings support mounting evidence that derepression and signaling dynamics are instrumental for the interpretation of graded morphogen-signaling(*39*).

The main principle of the mechanism that interprets graded NODAL-signaling is based on SNON, a SMAD2/3 target-gene repressor and Arkadia a signaling-activated derepressor (Fig. **6**). This principle is conserved because the interpretation of the TGFβ morphogen signaling, DPP-MAD, in *Drosophila*, is controlled by Brinker, a MAD target-gene repressor, and Schnurri, a derepressor, which complexes with activated MAD effectors to suppress Brinker leading to derepression(*40–42*). Furthermore, Arkadia and SNON are expressed broadly and involved in cancer(*43–45*), suggesting that signaling-dependent derepression and graded transcriptional responses are intrinsic to the TGFβ-SMAD2/3 signaling pathway.

**Fig. 6:**
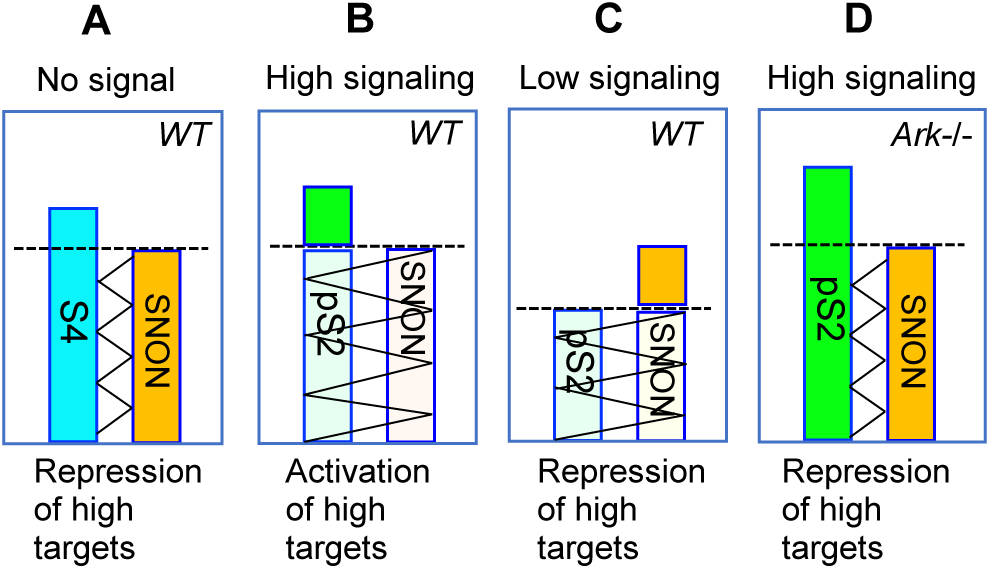
Signaling-induced derepression controls the activation of high-signaling responses. Model showing that absence of signaling (**A**), SNON establish repression via SMAD4 complex formation in a subset of targets. Under high signaling (**B**), high levels of SMAD2 saturate SNON causing degradation and removal of SNON. This leads to de-repression and activation of repressed targets. Under low signaling (**C**), SMAD2 reduces SNON causing partial derepression and activation of weakly repressed targets, while highly-repressed targets remain repressed. Absence of Arkadia (**D**) leads to accumulation of SMAD2-SNON complexes and maintenance of repression on targets.

## Supporting information

Auxiliary suppl Table

## Acknowledgments

We wish to thank Prof. Susumu Itoh (Showa Pharmaceutical Univ. Tokyo) for donating the Smad7 KO mice and Prof. Kunxin Luo (U.C. Berkeley) for the SnoN KO mice. We thank Dr. Vikas Sharma, Michael Normal and Nabiha Husain for preliminary experiments. Drs. Tristan Rodriguez and Aida DiGregorio (ICL) for help with in-situ hybridizations, Eleni Ioannou (ICL), Zoe Webster (MRC/ICL) and Dr. Christophe Galichet (Crick Institute) for technical assistance and animal procedures. We also thank Prof. Kunxin Luo for critical reading of the MS and suggestions.

## Authors contributions

J.M.C. and M.I. contributed equally by designing, conducting and analyzing experiments; and by providing a draft of figures, methods and results. VE raised funding; designed and conducted experiments; interpreted the combined results; and wrote the manuscript.

## Data and materials availability

The RNA sequencing data have been deposited in NCBI Gene expression Omnibus (accession number GSE118005).

## List of Supplementary Materials

Materials and Methods

Fig S1 – S5

## Other Supplementary Materials for this manuscript include the following

Caption: **Table S** (see **Table S** containing the list of Activin-induced genes in the Auxiliary supplementary data).

