## Supplemental materials for "Arkadia via SNON enables NODAL-SMAD2/3 signaling effectors to transcribe different genes depending on their levels"

**This PDF file includes:**

Materials and Methods

Fig S1 – S5

**Other Supplementary Materials for this manuscript include the following:**

Caption: **Table S** (see **Table S** containing the list of Activin-induced genes in the Auxiliary supplementary data).

**Materials and Methods**

Mice and Embryo methods

**Ethical compliance:** All procedures including breeding of genetically modified mice and harvesting mouse embryos for experiments were covered by a UK Home Office license under the name of Tristan Rodriguez. All laboratory mice were housed in the Central Biological Services of Imperial College London at the Hammersmith Hospital Campus.

Mice were backcrossed >5 generations to 129Sv/Ev background before they were used for WISH, X-gal staining, RNA or protein extraction. Mice were maintained in pathogen-free environment under a Home Office license (Scientific Procedures) Act 1986. *SnoN*-/- (homozygous for the *Skil*$^{tm2Spw}$ allele(*1*)) mice were kindly donated by Kuxin Luo (Univ. of California Berkeley). *Smad7*-/- mice were kindly donated by Susumu Itoh (Showa Pharmaceutical Univ.). *Ark*-/- mice were generated by VE(*2*). Mouse genotyping was performed as in the mouse strain references and by Transnetyx, Inc automatic genotyping.

**Whole-mount *in-situ* hybridization** (WISH) and X-gal staining was carried out as described(*2, 3*). SnoN riboprobe was generated in the laboratory by PCR. T7RNA polymerase promoter was introduced at the 5`-end of the primers designed for the gene of interest, enabling direct in vitro transcription of purified PCR fragments. More detailed info about the primers used can be found in the list of primers. For *Shh*, *Six3*, *Cel*, *Foxa2* probes see references(*2-4*).

**QPCR:** Total RNA was extracted from individual whole embryos at E 8.5 stage, using RNeasy Micro kit (Qiagen, 74134). cDNA was generated using the iScript reverse transcription supermix (BioRad, 1708840) and QPCR experiments were performed using QuantiFast SYBR Green PCR Kit (Qiagen, 204054), MicroAmp Optical 96 well reaction plates (Applied Biosystems, 4306737) and Stratagene mx3005 qPCR system and software. QPCR experiments were normalized against β-Actin. All p values were determined using one-way ANOVA method (Prism_GraphPad software). Primers/oligonucleotide-sequences used can be found in below).

**Western blots:** Total protein extraction was harvested from individual embryos lysed by adding 10 µl of NP-40 buffer (150Mm NaCl, 50mM Tris pH 8, 20mM NaF, 1%NP-40, 1mM EDTA pH 8, plus protease and phosphatase inhibitors). After centrifugation at 100,000 g for 10 min, equal amount of protein extracts was boiled in 2× Laemelli buffer followed by standard SDS-polyacrylamide gel electrophoresis (SDS-PAGE) and transferred onto nitrocellulose membrane. For analysis with antibodies see Cell culture methods.

Cell culture methods

Mouse embryonic fibroblasts (MEFs) were cultured in DMEM containing 10% FBS and supplemented with 1X Penicillin/Streptomycin and L-Glutamine. Mouse embryonic stem cells were cultured on inactivated MEF feeders and maintained in ES medium containing 15%FBS/DMEM with 1X Pen/Strep and L-Glutamine, 50 µM β-mercaptoethanol and supplemented with additional LIF made in house. Cells were maintained in a humidified incubator at 37 °C and 5% CO2. Prior to performing experiments, cells were maintained in full growth medium supplemented with 10µM SB431542 (Sigma Aldrich, S4317) to inhibit endogenous Smad2/3 signaling. Cells were then washed 2 times in PBS and replaced with medium containing fresh SB431542 or 5ng/mL TGFβ1 (Peprotech, 100-21C) or 10ng/mL Activin A (Sigma Aldrich, A4941) for the indicated time. In certain experiments the concentration of ligand was adjusted; exact experimental conditions are further outlined in the text.

**Western blots:** Cells were washed once in ice-cold PBS and proteins were collected in lysis buffer (20 mM Tris-HCl pH 8.0, 1% Triton-X, 150 mM NaCl, 2 mM EDTA) supplemented with protease inhibitor cocktail (Roche Diagnostics) and cleared by centrifugation at 12,000x g at 4 °C for 10 min. Equal amounts of protein from each sample were separated by standard SDS-polyacrylamide gel electrophoresis (SDS-PAGE) and transferred to nitrocellulose membrane. Membranes were blocked for 1 h in 5% milk/TBS-T and incubated overnight at 4 °C with primary antibody in TBS-T. Following 3 washes in TBS-T, corresponding HRP-conjugated secondary antibody (Santa Cruz Biotechnology) was added in TBS-T at a concentration of 1:5000 and then incubated for a further 1 h at room temperature. Membranes were washed 3 times in TBS-T and antibody binding was visualized with the enhanced chemiluminescence detection system (Thermo Fischer Scientific Inc.). Images were captured using the GeneGnome XPQ NPC System (Synoptics Limited) and band intensities were calculated. Antibodies used were c-Ski (Santa Cruz Biotechnology, H-329: SC-9140), SnoN (Santa Cruz Biotechnology, H-317: SC-9141), pSmad2 (phosphorylated at Ser465/467 Cell Signaling Technologies: 3101), Smad2/3 (BD Biosciences: 610842); β-Actin (Santa Cruz Biotechnology, SC-47778); β-Tubulin Merck Millipore: (05-661); PCNA (MAB424R).

**Proximity Ligation assay (PLA)** was performed using the Duolink *in-situ* PLA kit (Sigma) as described(*5*). Cells were cultured overnight on 8-well chamber slides, stimulated with TGFβ (or SB43152 control) for the indicated timepoints Primary antibodies were used at a concentration of 1:500 in blocking solution and left overnight at 4C in a humidified chamber. Dots per nuclei were counted and plotted in bar graphs; each experiment was repeated at least three independent times.

**QPCR** were performed as described in mice and embryo methods. Primers sequences are shown in Extended data table 2 and predesigned quantitech primer assays (Qiagen, 249900) were used for other genes including Gal (QT00109970), *RhoB* (QT00249648), Plekha2 (QT00152103), Ncor2 (QT00166439), FoxA2 (QT00242809) and Sox17 (QT00160720). Expression levels of the target genes were quantified by normalizing against β-actin using the delta Ct method.

**Embryoid body (EB) differentiation.** Mouse *WT* and *Ark*-/- ESCs were trypsinized, and pre-plated on 2% gelatin coated plates for ½ hour to remove feeders, and then transferred to non-coated bacterial dishes at a concentration of 1 X $10^6$ cells per 10 cm dish in ES growth media without LIF for 48 h to form EBs in suspension(*6*). The EBs were collected and allowed to sediment in falcon tubes where they were washed 2 times with PBS and then resuspended in serum replacement medium (2% serum replacement/DMEM supplemented with 1X Pen/Strep and L-Glutamine and 0.45mM Monothioglycerol) containing FGF 10ng/ml (Catalogue) and 50ng/ml A. The EBs were then transferred to new bacterial dishes and maintained for an additional 48h prior to imagining under an inverted fluorescent microscope using the Infinity3 camera (Luminera, Canada). EB's were then centrifuged, washed in PBS and RNA or for protein extraction for further analysis.

**RNA sequencing**: ESCs were separated from feeders by pre-plating as above then cultured for one passage in 2i medium consisting of N2B27 supplemented with Glutamine and Pen/Strep, 50µM β-mercaptoethanol, 3µM CHIRON (Cayman Chemical, 13122), 1µM PD03 (Sigma Aldrich, PZ0162) and LIF (homemade). Cells were trypsinized and plated into Cellbind dishes (Corning, 734-4057) coated with 40µg/dish fibronectin (Millipore, FC010) at a concentration of 3X$10^6$ cells per dish in N2B27 medium without 2i. 24 h later, cells were switched to fresh

N2B27 medium containing 10μM SB43152 for 4 h to inhibit endogenous TGFβ signaling. Cells were washed 2 times in PBS and then treated with 10ng/mL Activin A for 3 or 6 h or SB431542 for 3 hours prior to harvesting for RNA as described for qPCR (see mouse and embryo methods). The integrity of the RNA was checked using Agilent 2100 Bioanalyzer and a total of 1 μg RNA with a RIN>8 was used for sequencing. RNA sequencing was performed by GATC Biotech (Constance, Germany). In ESCs treated in parallel were processed for western blot analysis.

**CRISPR-Cas9:** CRISPR guide RNAs (sgRNA) were designed using the Feng Zhang lab Target Finder design tool (http://crispr.mit.edu/). We designed a total of 4 guide-RNA primer pairs (see table below). Each guide oligonucleotide was purchased with its complementary oligonucleotide (Sigma Aldrich) and annealed using the T4 Polynucleotide Kinase Kit (New England Biolab, M0201). Annealed guides were then cloned into the pSpCas9(BB)-2A-Puro (PX459) V2.0 (Addgene, plasmid 62988, PMID: 24157548). To create Ark/SnoN double null ESCs, the *Ark*-/- ESC-line 39 (A-/- 39) was used. ESCs were weaned from feeders and cultured in 2i medium plus LIF on gelatinized Cellbind dishes (as described for RNA sequencing) for transfection and puromycin selection. Transfection was performed by nucleofection according to manufacturer's instructions for the Nucleofector™ 2b Device (Lonza) and the Mouse ES cell Nucleofector kit (Lonza, Cat No. VPH-1001). 36 hours after transfection the cells were selected by 2μg/ml puromycin for 48 hours. Colonies were transferred onto 96 well plates with feeders on DMEM-LIF medium for expansion.

**Primers used in Methods**

*In-situ* probe:

*SnoN* Forward: 5'AAGCTGAACGGCATGGAG3'

*SnoN* Reverse: 5'GAGTAATACGACTCACTATAGGGATGACGAACGTCTGGGGA3'

Quantitative RT-PCR:

*β-Actin* Forward: 5'-TGGCTCCTAGCACCATGA-3'

*β-Actin* Reverse: 5'-CCACCGATCCACACAGAG-3'

*Smad7* Forward: 5'-GGCCGGATCTCAGGCATTC-3'

*Smad7* Reverse: 5'-TTGGGTATCTGGAGTAAGGAGG-3'

*Lefty1* Forward: 5'-TGTGTGTGCTCTTTGCTTCC-3'

*Lefty1* Reverse: 5'-GGGGATTCTGTCCTTGGTTT-3'

*Lefty2* Forward: 5'-GATGGCTCCAATCGCACTG-3'

*Lefty2* Reverse: 5'-GTGGATGGACACGAGCCTAGAG-3'

*Cxcr4* Forward: 5'- TTTCAGCCAGCAGTTTCCTT-3'

*Cxcr4* Reverse: 5'- TCAGTGGCTGACCTCCTCTT-3'

CRISPR/Cas9 guide RNA sequences

*SnoN exon1* Guide 1: 5'-TGACCGACATTCATGCCAAT-3' *SnoN exon1* Guide 2: 5'-TAGAGAAACACGGTTTCGCT-3' *SnoN exon1* Guide 3: 5'-GCGAATGCATGACGAACGTC-3' *SnoN exon1* Guide 4: 5'-CGGGTGTCCCTAGGTATTTT-3'

Fig. S1

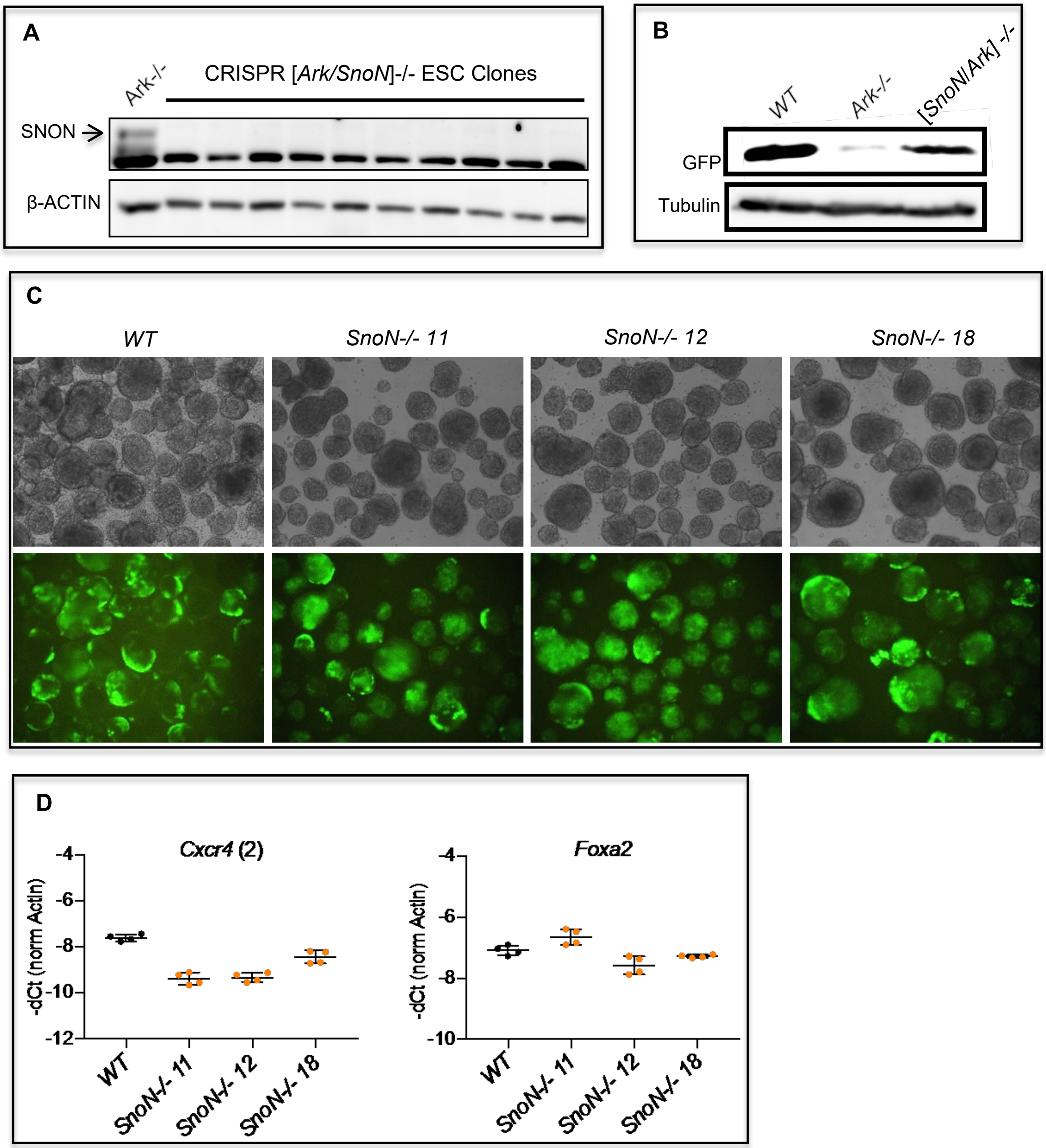


**Fig. S1:** (**A**) Western blot analysis showing loss of SNON protein in 10 *SnoN*-CRISPR clones generated in an *Ark*-/- ESC-line. (**B**) Western blot analysis with GFP antibody on ADE-ME differentiated EBs carrying *Hex-Gfp* transgene showing rescued of GFP expression in [*SnoN/Ark*]-/- EBs. (**C**) Bright field (top) and fluorescence (bottom) images of EBs differentiated towards ADE-ME from 10 *SnoN*-/- and one *WT* ESC-lines. All lines show successful ADE-ME differentiation. (**D**) QPCR on the above differentiated EBs with definitive endoderm (*Cxcr4*) and ME (*Foxa2*) markers. Note that *Cxcr4* expression is reduced in the three *SnoN-/-* EBs compared to *WT* suggesting that *SnoN*-/- ESCs may exhibit a mild phenotype.

**Fig. S2**

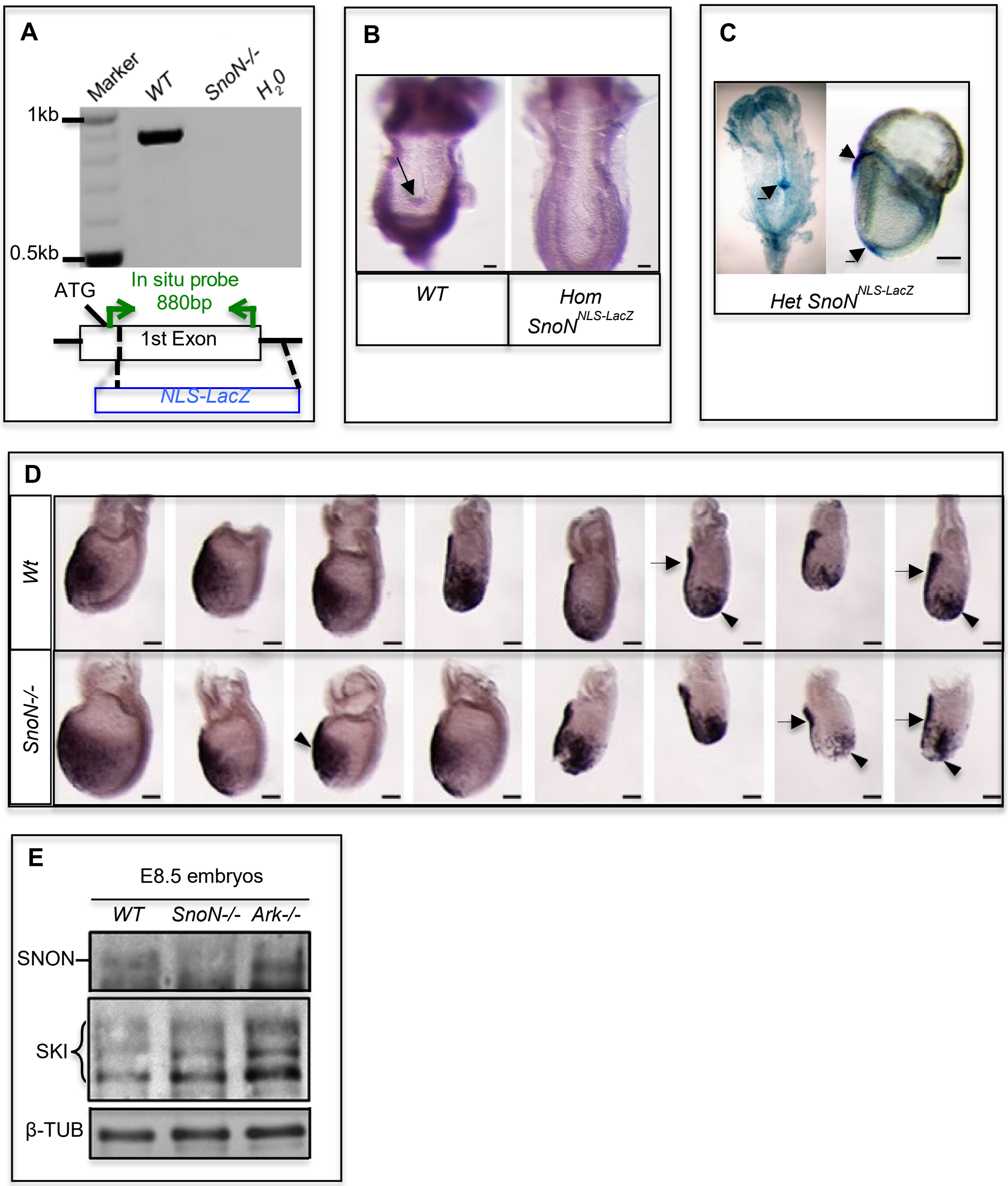


**Fig. S2:** (**A**) Agarose gel and schematic representation show the *in-situ* riboprobe generated from the first coding exon of *SnoN* gene containing all the conserved domains interacting with SMAD2/3, SMAD4 and histone deacetylases. Green arrows indicate the position of the primers used; the blue box indicates the targeted replacement of the endogenous *SnoN*-exon with *β-galactosidase* gene (*LacZ*) containing nuclear localization signal (NLS) which is expressed by the endogenous promoter of *Skil*$^{tm2Spw}$ allele. (**B**) Images of WISH stained E8.5 mouse embryos with the above probe showing loss of expression in the embryos homozygous for the *Skil*$^{tm2Spw}$ allele embryos. (**C**) Images of a 2-somite E8.5 embryos (right panel) and late head-fold E7.5 (left panel) heterozygous for the *SnoN*$^{\text{NLS-lacZ}}$ allele stained with X-gal. Limiting the X-gal staining to <30 minutes allows quantification of expression levels differences. Arrows and arrowhead indicate high levels of *SnoN*-expression in the node-ME and ADE respectively. (**D**) Images of WISH with the ADE marker (*Cer)* probe on *WT* and *SnoN*-/- embryos stage E7-E7.5 with anterior to the left and scale bar 100µm. Arrowheads and arrows indicate ADE and AVE respectively. (**E**) Western blot analysis of E8.5 embryo-extracts with antibodies and genotypes as indicated. SKI upregulation in *SnoN*-/- embryos may reflect expansion of *Ski* expression-pattern in cells that normally express high *SnoN* and low *Ski*.

**Fig. S3**

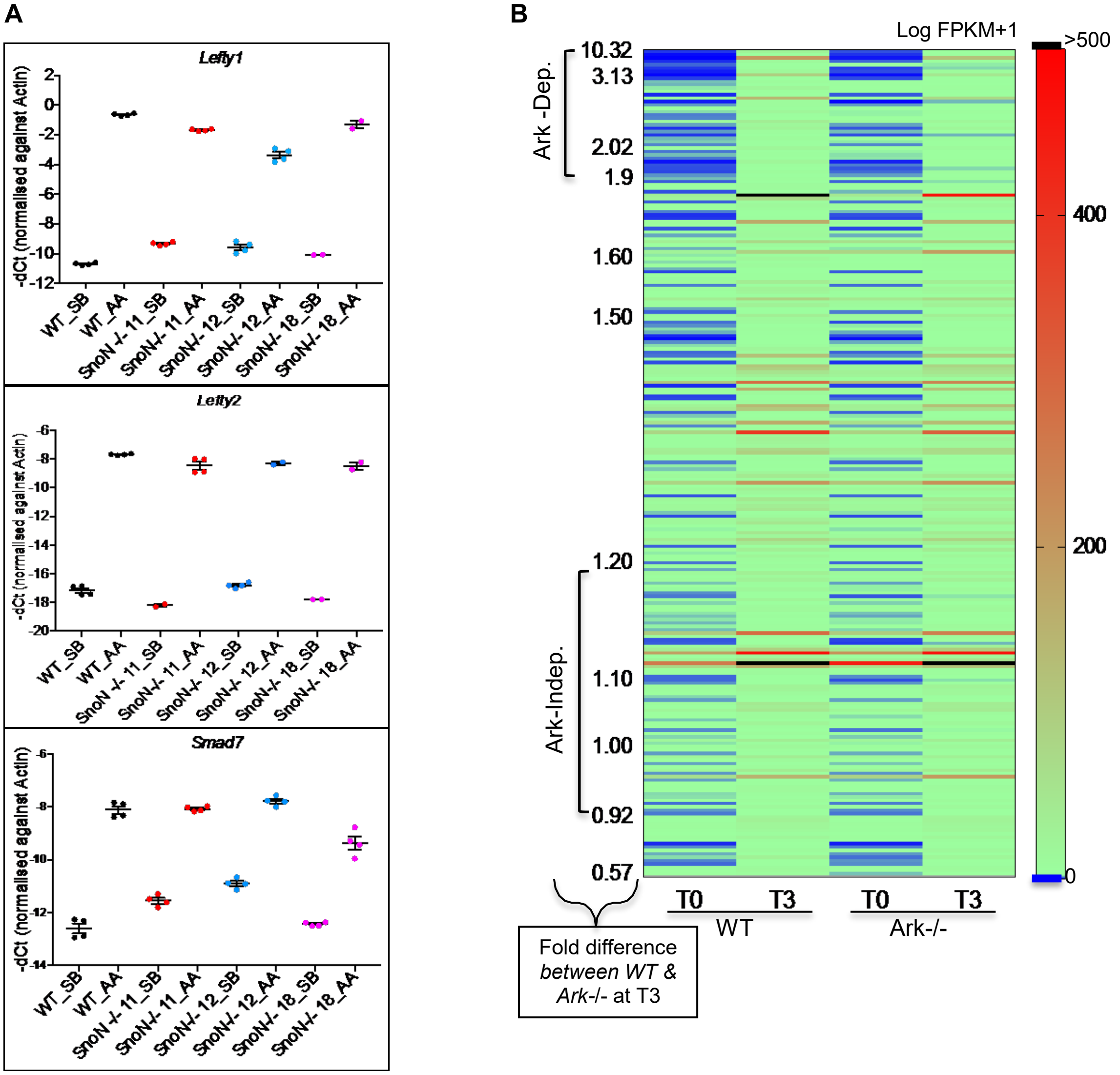


**Fig. S3:** (**A**) Signaling induction of known SMAD2/3 target genes assayed via qPCR in *WT* and *SnoN*-/- ESCs lines treated with the SB inhibitor (4hr) or Activin (3hr) showing that in the absence of SNON there is signaling induction comparable to *WT*. (**B**) Heat-map of all SMAD2/3 genes with 2-fold Activin induction at T3 in both *WT* ESC-lines identified by the RNA-sequencing experiment. Arkadia-dependent targets (1.9 to 10.32-fold difference between *WT* and *Ark*-/- ESCs) are shown at the top and Arkadia-independent at the bottom.

Fig. S4

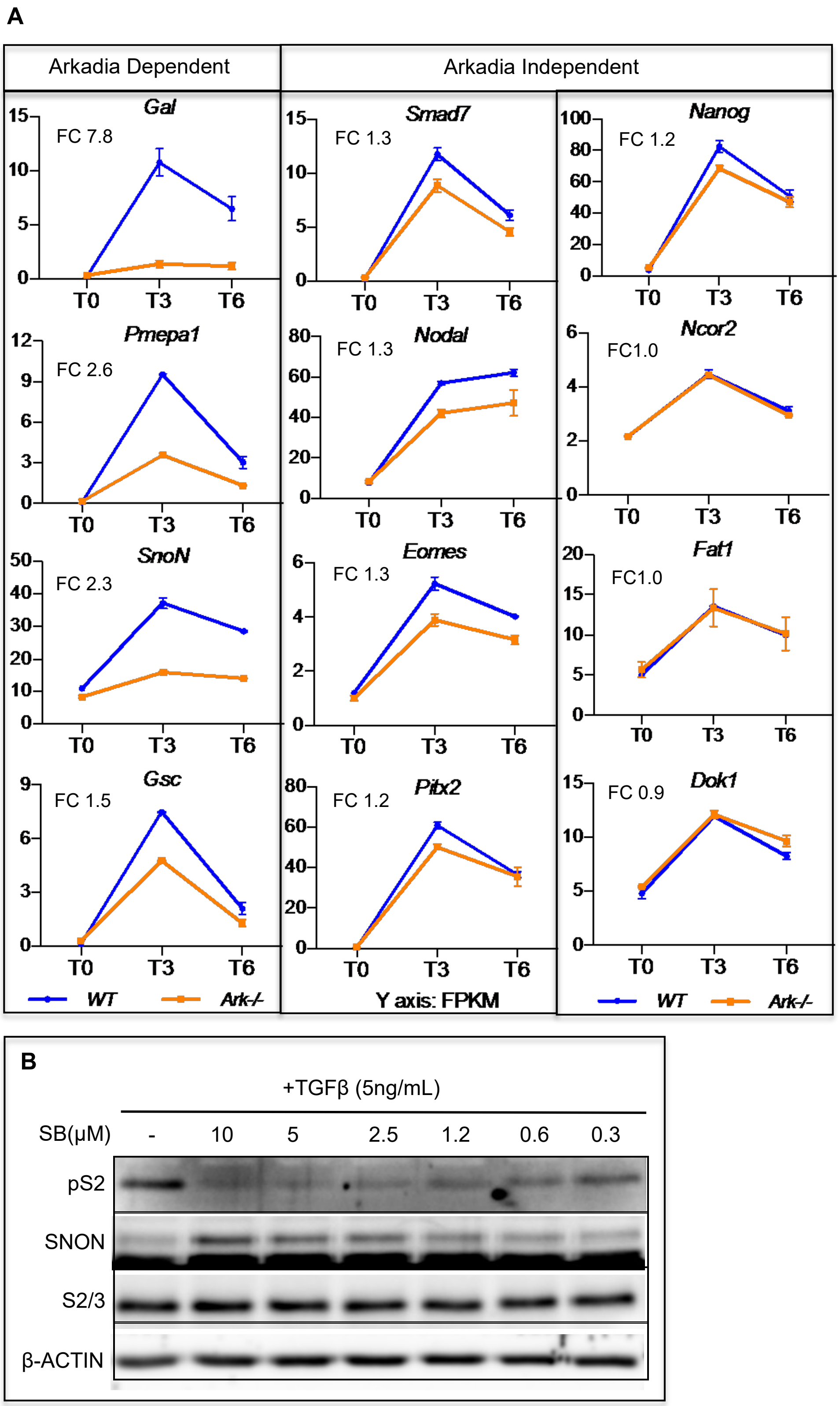
A
Arkadia Dependent
Arkadia Independent
Gal
FC 7.8
Smad7
FC 1.3
Nanog
FC 1.2
Pmepa1
FC 2.6
Nodal
FC 1.3
Ncor2
FC1.0
SnoN
FC 2.3
Eomes
FC 1.3
Fat1
FC1.0
Gsc
FC 1.5
Pitx2
FC 1.2
Dok1
FC 0.9
T0 T3 T6
WT
Ark-/-
Y axis: FPKM
B
+TGFβ (5ng/mL)
SB(μM) - 10 5 2.5 1.2 0.6 0.3
pS2
SNON
S2/3
β-ACTIN

**Fig. S4**: (**A**) Graphs of additional Arkadia-dependent and -independent SMAD2/3 targets from the RNA-sequencing experiment. (**B**) Western blot analysis of proteins (as indicated) derived from *WT*-ESCs treated for 1hour with TGFβ ligand titrated with different amounts of SB inhibitor as indicated.

**Fig. S5**

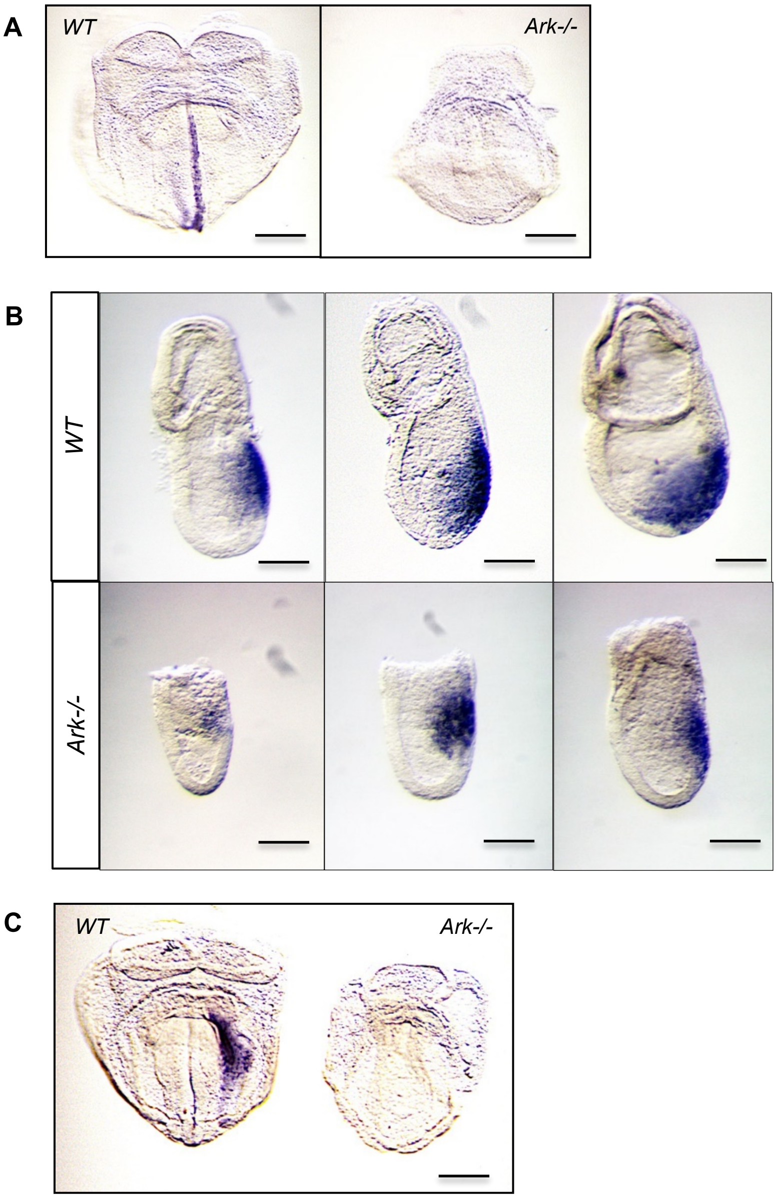


**Fig. S5**: (**A**) Expression analysis in early somite-stage embryos using WISH for *Lefty1* showing expression in the midline mesendoderm of *WT* and its absence in *Ark*-/- embryos, reflecting the failure of *Ark*-/- embryos to develop midline mesendoderm (Fig. 2). (**B**) WISH with *Lefty2* on three *WT* and three *Ark*-/- E7.5 embryos showing *Lefty2* expression in the anterior-PS (APS) and the more posterior-PS, while *Ark*-/- embryos exhibit reduced *Lefty2* expression in the APS domain. This most likely reflects loss of APS in Ark-/- embryos. (**C**) WISH with *Lefty2* expression in 4-somite stage embryos shows lack of expression in the left lateral mesoderm in the *Ark*-/- embryo. However, this reflects the failure of *Ark*-/- embryos to form node, which is essential for the induction of left lateral plate mesoderm(*7*). The scale bars are 0.1mm. The data of this figure are from *R. Andrew, thesis, Imperial College London, (2003)*.

**Other Supplementary Materials for this manuscript include the following:**

Caption: **Table S:** Activin induced targets in *WT* and *Ark-/-* ESCs derived from the RNA-sequencing results containing the expression profile of genes (logFPKM+1) at T0; T3 and T6 time points in two different *WT* (WT-18 and WT-26) and two different *Ark*-/- (A-/- 25 and A-/-39) ESC-lines. The genes on this list are all induced ≥2-fold by Activin in both *WT* ESC lines at T3. 52 genes in red font are considered Arkadia dependent because they exhibit Activin-induction difference ≥1.7-fold between *WT* and *Ark*-/- ESCs at T3 (see last column). The values of the last column show the induction difference of each gene between ESCs *WT* and *Ark*-/- genotype at T3. The induction value was averaged between the two ESC lines with the same genotype. (see **Table S** containing the list of Activin-induced genes in the Auxiliary supplementary data)
