## Supplementary material for "Arkadia via SNON enables NODAL-SMAD2/3 signaling effectors to transcribe different genes depending on their levels": Auxiliary suppl Table

| 0 | gene symbol | refseq_ID | WT18 @T0 | WT18 @T3 | WT18 @T6 | WT26 @T0 | WT26 @T3 | WT26 @T6 | A-/-25 @T0 | A-/-25 @T3 | A-/-25 @T0 | A-/-39 @T0 | A-/-39 @T3 | A-/-25 @T6 | Induction Difference WT/A-/-@T3 |
|---|---|---|---|---|---|---|---|---|---|---|---|---|---|---|---|
| 1 | Gal | NM 001329667 | 0.301309 | 9.89265 | 5.69885 | 0.185166 | 11.6717 | 7.28585 | 0.48687 | 1.59483 | 1.37061 | 0.190167 | 1.14273 | 0.949186 | 10.32508 |
| 2 | Ctla2b | NM 007797 | 0.0318262 | 2.02704 | 1.78561 | 0.114925 | 1.57846 | 1.97079 | 0.164566 | 1.10102 | 1.64303 | 0.282756 | 1.1325 | 1.45263 | 7.23897088 |
| 3 | Lefty1 | NM 010094 | 0.0209895 | 93.7814 | 40.8536 | 0.0515953 | 111.231 | 40.7898 | 0.108531 | 70.4708 | 35.4317 | 0.127173 | 49.3926 | 27.7864 | 6.38317954 |
| 4 | Col8a2 | NM 199473 | 0.30886 | 6.06304 | 2.31302 | 0.0904035 | 7.58336 | 0.767249 | 0.238712 | 3.02681 | 2.57071 | 0.587088 | 2.49049 | 1.9739 | 6.11716687 |
| 5 | Exoc3l4 | NM 001289489 | 0.463469 | 1.37623 | 1.17095 | 0.379759 | 1.22591 | 1.25135 | 1.02706 | 0.811566 | 1.39903 | 1.3372 | 1.0446 | 1.96305 | 3.94403895 |
| 6 | Fam101a | NM 028443 | 0.369192 | 2.07987 | 2.14524 | 0.108821 | 2.31413 | 1.06605 | 0.378268 | 1.0108 | 0.895706 | 0.163632 | 0.680192 | 0.377045 | 3.93893244 |
| 7 | Gsc | NM 010351 | 0.184819 | 7.47953 | 2.42119 | 0.100959 | 7.44442 | 1.78077 | 0.292003 | 4.77111 | 1.48255 | 0.342161 | 4.74768 | 1.09594 | 3.77981852 |
| 8 | Lefty2 | NM 177099 | 0.0283309 | 47.8244 | 17.3232 | 0.0696418 | 49.2998 | 21.7855 | 0.0651072 | 27.8549 | 19.4114 | 0.0858272 | 21.008 | 12.9917 | 3.56223964 |
| 9 | Adap2 | NM 172133 | 0.452372 | 1.38907 | 1.11117 | 0.0926667 | 1.76335 | 0.75439 | 0.292386 | 1.00817 | 1.46036 | 0.34261 | 1.23527 | 0.838274 | 3.13311604 |
| 10 | Tspear | NM 001287074 | 0.675389 | 4.16243 | 3.04004 | 0.857531 | 5.07691 | 3.88416 | 1.12046 | 2.83146 | 2.81364 | 1.30594 | 1.99419 | 2.59179 | 2.98056093 |
| 11 | Prkg2 | NM 008926 | 0.690902 | 1.44866 | 3.38082 | 0.744901 | 1.65969 | 3.12072 | 2.29086 | 1.37017 | 4.76059 | 2.03351 | 1.92633 | 3.61747 | 2.79852811 |
| 12 | Nfic | NM 008688 | 1.01057 | 11.6167 | 2.97874 | 0.828067 | 8.20672 | 4.49403 | 0.725747 | 4.31671 | 1.77947 | 1.70083 | 3.27053 | 2.24718 | 2.71963934 |
| 13 | Utrn | NM 011682 | 1.41657 | 2.89986 | 3.94349 | 0.773813 | 2.99099 | 1.83736 | 1.62771 | 2.36777 | 2.49438 | 2.38413 | 1.71918 | 2.8 | 2.71738204 |
| 14 | Zdhhc1 | NM 175160 | 0.208425 | 1.33908 | 1.02392 | 0.170781 | 1.50303 | 0.834185 | 0.431083 | 1.16126 | 1.17442 | 0.29466 | 0.860027 | 0.617961 | 2.71280126 |
| 15 | Cd40 | NM 170704 | 1.88597 | 72.1271 | 30.8038 | 1.41979 | 71.523 | 26.7269 | 2.1088 | 40.4648 | 27.9824 | 2.69464 | 37.2064 | 30.378 | 2.68576553 |
| 16 | Zbp1 | NM 021394 | 0.112214 | 4.78397 | 4.83971 | 0.164604 | 4.0169 | 6.18825 | 0.03325 | 0.59483 | 1.75392 | 0.089545 | 0.870816 | 1.70653 | 2.42756241 |
| 17 | Krt17 | NM 010663 | 0.556111 | 5.08607 | 3.49692 | 0.692617 | 4.33546 | 3.31387 | 0.632609 | 1.90863 | 2.14125 | 0.628959 | 2.1867 | 1.91262 | 2.37232098 |
| 18 | Sox15 | NM 009235 | 0.94776 | 2.96427 | 2.02295 | 1.28537 | 3.12112 | 2.36169 | 1.71803 | 1.74804 | 2.4937 | 1.15508 | 1.66584 | 1.93794 | 2.2587925 |
| 19 | Itpkb | NM 001081175 | 0.954906 | 8.0553 | 2.11005 | 0.774924 | 9.5778 | 1.71378 | 0.601228 | 3.93901 | 1.93867 | 1.29804 | 3.46279 | 0.891354 | 2.2556301 |
| 20 | Gbp2b | NM 010259 | 0.488584 | 2.43029 | 4.36277 | 0.529859 | 2.28261 | 5.06441 | 0.990717 | 1.7454 | 4.25101 | 0.914205 | 2.17998 | 4.49493 | 2.23863852 |
| 21 | Ddx42 | NM 028074 | 0.404225 | 1.47311 | 1.41843 | 0.662432 | 1.68823 | 1.7334 | 1.1446 | 1.99478 | 1.69472 | 1.69109 | 1.75206 | 1.88334 | 2.22857067 |

| | | | | | | | | | | | | | | |
|---|---|---|---|---|---|---|---|---|---|---|---|---|---|---|
| 22 | Grm6 | NM 173372 | 6.87761 | 15.5076 | 19.9037 | 3.223 | 20.1141 | 19.9663 | 4.87279 | 10.4236 | 18.6602 | 5.01104 | 8.47481 | 11.5333 | 2.21795577 |
| 23 | Mkx | NM 177595 | 0.635827 | 8.30334 | 3.87292 | 0.672707 | 8.1071 | 4.25256 | 0.659279 | 3.50348 | 3.35132 | 0.641154 | 3.95242 | 3.81613 | 2.18758983 |
| 24 | Smad7 | NM 001042660 | 0.197742 | 11.3622 | 5.77114 | 0.235427 | 12.1969 | 6.49326 | 0.383227 | 8.46444 | 4.83225 | 0.330212 | 9.30182 | 4.35161 | 2.17419164 |
| 25 | Zfp819 | NM 028913 | 0.592375 | 3.37381 | 1.27725 | 0.647129 | 3.64 | 1.32712 | 0.939919 | 2.06494 | 1.00746 | 0.566998 | 1.76449 | 0.71668 | 2.13230555 |
| 26 | Il15ra | NM 001271498 | 0.297189 | 1.11121 | 1.29561 | 0.1834 | 1.02077 | 1.07373 | 0.340362 | 0.7824 | 1.92293 | 0.310207 | 0.641509 | 1.54266 | 2.13085687 |
| 27 | Has2 | NM 008216 | 0.776619 | 5.17991 | 5.60576 | 0.680453 | 4.2187 | 5.33434 | 1.21928 | 3.26453 | 5.30757 | 1.07154 | 3.6581 | 5.08396 | 2.11279533 |
| 28 | Tnfrsf13c | NM 028075 | 0.861422 | 2.67665 | 0.97073 | 0.75469 | 2.27261 | 0.42307 | 0.774268 | 1.19583 | 1.04154 | 1.05051 | 1.47864 | 1.22658 | 2.07267643 |
| 29 | H2-M3 | NM 013819 | 0.613734 | 2.4161 | 3.4673 | 0.460977 | 2.61663 | 4.60567 | 0.661134 | 1.33929 | 2.58166 | 0.413173 | 1.11726 | 1.85757 | 2.03241206 |
| 30 | Slc6a15 | NM 175328 | 1.08967 | 9.90436 | 11.5052 | 1.38642 | 9.55098 | 8.46426 | 1.24448 | 4.4528 | 7.40684 | 1.49691 | 6.46533 | 9.29688 | 2.02329395 |
| 31 | Dync1h1 | NM 030238 | 0.500346 | 1.01634 | 0.506883 | 0.450118 | 1.32242 | 0.942418 | 0.896406 | 1.20485 | 1.02948 | 1.00975 | 1.19225 | 0.834455 | 1.96814049 |
| 32 | Mfsd4a | NM 001114662 | 0.784839 | 2.3576 | 1.55394 | 0.722833 | 3.07853 | 1.59475 | 0.989355 | 1.66945 | 1.90849 | 0.773316 | 1.55672 | 1.05777 | 1.96270493 |
| 33 | Cxcl16 | NM 023158 | 0.539064 | 1.37123 | 1.22226 | 0.407724 | 1.3335 | 1.33691 | 0.643234 | 1.24758 | 1.11958 | 1.17246 | 1.20779 | 1.29091 | 1.95789652 |
| 34 | Pmepa1 | NM 022995 | 0.0911098 | 9.49651 | 2.59601 | 0.117313 | 9.58251 | 3.46917 | 0.0448669 | 3.71286 | 1.30633 | 0.279257 | 3.43165 | 1.34581 | 1.9561478 |
| 35 | Slco4c1 | NM 172658 | 0.399184 | 3.03895 | 0.952458 | 0.284222 | 2.60152 | 1.25581 | 0.646413 | 2.06629 | 1.56645 | 0.368978 | 2.01992 | 0.950896 | 1.93359499 |
| 36 | Erap1 | NM 030711 | 0.362278 | 1.66773 | 2.3023 | 0.290529 | 1.4347 | 3.06786 | 0.438417 | 0.86751 | 2.25307 | 0.280214 | 0.832562 | 1.63403 | 1.92765347 |
| 37 | Wnt3 | NM 009521 | 0.310701 | 4.61589 | 2.08427 | 0.289699 | 4.57302 | 2.6681 | 0.480118 | 3.95168 | 2.41478 | 0.400304 | 3.0816 | 2.08064 | 1.92367321 |
| 38 | Mab21l3 | NM 172295 | 0.657859 | 2.40489 | 1.35251 | 0.536723 | 2.30065 | 1.13584 | 0.746167 | 1.44699 | 1.22994 | 0.521663 | 1.15921 | 1.03967 | 1.90852986 |
| 39 | Ptges | NM 022415 | 0.824138 | 3.64136 | 3.40544 | 0.695148 | 3.70857 | 3.47886 | 1.44135 | 3.13325 | 3.03466 | 0.905661 | 2.7064 | 2.67001 | 1.8893934 |
| 40 | Nxph4 | NM 183297 | 0.397603 | 1.52351 | 0.412359 | 0.361989 | 1.48529 | 0.270134 | 0.418793 | 1.00918 | 0.466747 | 0.401507 | 0.750616 | 0.16373 | 1.85427518 |
| 41 | Ebf2 | NM 010095 | 0.897553 | 5.30375 | 0.905487 | 1.29698 | 5.00714 | 0.956746 | 1.51959 | 2.86083 | 1.39221 | 0.572999 | 2.03962 | 0.3951 | 1.7951863 |
| 42 | Steap2 | NM 001103157 | 1.23014 | 5.443 | 4.29528 | 1.8203 | 4.41256 | 4.1934 | 1.92623 | 3.67012 | 3.90905 | 1.90959 | 3.68008 | 2.30879 | 1.78702975 |
| 43 | Sphk1 | NM 001172475 | 0.30765 | 1.44303 | 0.842072 | 0.351126 | 1.59004 | 1.19104 | 0.771699 | 2.24069 | 0.941559 | 0.68789 | 1.5602 | 0.952899 | 1.78257422 |
| 44 | Tdgf1 | NM 011562 | 13.875 | 280.108 | 225.288 | 16.5579 | 300.428 | 257.23 | 19.7601 | 210.065 | 213.589 | 23.4439 | 259.438 | 270.143 | 1.76669022 |

| | | | | | | | | | | | | | | | |
|---|---|---|---|---|---|---|---|---|---|---|---|---|---|---|---|
| 45 | Skil | NM 001039090 | 10.9271 | 35.495 | 28.5567 | 11.1067 | 38.6448 | 28.2133 | 8.80796 | 15.743 | 14.1159 | 7.96846 | 16.2048 | 14.0175 | 1.76074269 |
| 46 | Ctsw | NM 009985 | 0.397329 | 3.84549 | 3.94699 | 0.383046 | 3.87916 | 3.56018 | 0.502972 | 2.97913 | 3.76059 | 0.452463 | 2.41401 | 4.20972 | 1.75918733 |
| 47 | Serpine1 | NM 008871 | 1.81651 | 7.38099 | 7.02959 | 1.76237 | 7.14596 | 7.57009 | 3.92802 | 9.25159 | 13.2654 | 2.86968 | 6.55957 | 10.1933 | 1.74915968 |
| 48 | Col2a1 | NM 001113515 | 2.11376 | 4.49228 | 3.18655 | 1.42635 | 4.50128 | 2.96833 | 2.39149 | 3.84656 | 3.32119 | 2.45994 | 3.47416 | 3.75832 | 1.74827166 |
| 49 | Clic5 | NM 172621 | 0.60732 | 1.25044 | 1.26571 | 0.40996 | 1.08001 | 1.66667 | 0.861819 | 1.15364 | 1.17611 | 0.612661 | 0.824865 | 0.849881 | 1.7480145 |
| 50 | Ctla2a | NM 001145799 | 0.189621 | 1.49297 | 2.54621 | 0.233059 | 1.3551 | 2.42434 | 0.408533 | 1.54952 | 1.74367 | 0.255311 | 1.03558 | 1.78033 | 1.74388855 |
| 51 | Rin2 | NM 028724 | 0.141551 | 2.10639 | 2.12093 | 0.318957 | 2.6209 | 1.84909 | 0.27447 | 1.97166 | 1.20471 | 0.285881 | 1.76083 | 1.9935 | 1.73110824 |
| 52 | Nanog | NM 001289831 | 3.95282 | 79.9585 | 53.7743 | 3.44656 | 85.1311 | 48.1846 | 5.44098 | 69.8612 | 44.7091 | 4.97947 | 67.2092 | 49.4779 | 1.70590393 |
| 53 | Socs1 | NM 001271603 | 1.83886 | 5.04924 | 2.68185 | 1.22422 | 5.40392 | 2.87488 | 1.48566 | 3.71394 | 3.20422 | 1.68282 | 2.8942 | 2.17229 | 1.69680561 |
| 54 | Ppp4r4 | NM 028980 | 0.4102 | 3.31025 | 6.45651 | 0.299188 | 3.126 | 5.67563 | 0.282855 | 1.68905 | 4.01491 | 0.307769 | 1.53366 | 4.64102 | 1.69044473 |
| 55 | Pnpla3 | NM 054088 | 2.20438 | 6.91135 | 4.52046 | 2.66914 | 5.85959 | 5.01079 | 2.78754 | 3.91638 | 3.9392 | 2.91595 | 5.30967 | 3.49755 | 1.65245319 |
| 56 | Timp4 | NM 080639 | 0.504536 | 7.38873 | 4.80467 | 0.826818 | 6.35392 | 5.95439 | 0.618757 | 3.87062 | 3.64477 | 0.764234 | 5.58149 | 4.89044 | 1.6468493 |
| 57 | Parp9 | NM 030253 | 1.31365 | 3.4264 | 7.01782 | 1.16093 | 3.41509 | 9.01404 | 1.55396 | 2.67633 | 5.70603 | 1.54242 | 2.55182 | 6.16855 | 1.64361811 |
| 58 | Ccl25 | NR 033527 | 19.0452 | 38.4003 | 36.8865 | 13.9567 | 37.3995 | 32.26 | 31.8793 | 41.7931 | 50.7729 | 28.9287 | 44.8634 | 52.5105 | 1.64090532 |
| 59 | Fhl2 | NM 010212 | 1.30537 | 5.21965 | 5.61122 | 1.146 | 5.88343 | 5.36446 | 2.29008 | 6.20803 | 9.24901 | 1.55358 | 4.52637 | 7.63694 | 1.62374132 |
| 60 | Fgd6 | NM 053072 | 0.65928 | 1.39811 | 0.83561 | 0.725776 | 1.48345 | 0.754947 | 0.746161 | 1.0432 | 0.875189 | 1.04771 | 1.24864 | 0.859186 | 1.60803915 |
| 61 | Id1 | NM 010495 | 12.2322 | 41.0422 | 14.5241 | 11.3687 | 46.3757 | 10.1394 | 47.4557 | 103.891 | 21.869 | 35.9658 | 87.6554 | 23.0035 | 1.6069708 |
| 62 | Kcnmb4os2 | NR 130644 | 1.22189 | 3.1168 | 2.9148 | 0.601778 | 2.48494 | 1.93169 | 1.22354 | 2.47127 | 2.15582 | 1.38336 | 2.96582 | 2.09402 | 1.60437588 |
| 63 | Fam214a | NM 001113283 | 3.1502 | 8.14698 | 2.48564 | 2.56024 | 8.94318 | 4.45125 | 1.47461 | 4.31885 | 3.32032 | 3.3187 | 2.88918 | 3.95702 | 1.60007008 |
| 64 | Tagln3 | NM 019754 | 0.885067 | 2.91329 | 2.88932 | 0.842179 | 2.55968 | 2.34885 | 1.20562 | 2.79966 | 1.91038 | 1.23973 | 2.04432 | 2.11624 | 1.59422636 |
| 65 | Herc6 | NM 025992 | 2.45085 | 5.38754 | 8.78126 | 2.3744 | 4.953 | 11.0248 | 2.05158 | 3.2965 | 9.58023 | 3.33932 | 3.63183 | 8.1147 | 1.59004763 |
| 66 | Sptan1 | NM 001177668 | 0.648165 | 2.14338 | 1.91052 | 0.763451 | 1.65805 | 1.93661 | 1.43136 | 2.70848 | 1.56932 | 1.79995 | 2.80235 | 2.6424 | 1.58840048 |
| 67 | Lgr4 | NM 172671 | 0.274688 | 1.55717 | 1.02557 | 0.315105 | 1.87379 | 0.916156 | 0.331411 | 1.34679 | 0.709403 | 0.443815 | 1.46681 | 0.977305 | 1.57629614 |

| | | | | | | | | | | | | | | | |
|---|---|---|---|---|---|---|---|---|---|---|---|---|---|---|---|
| 68 | Trim34a | NM 030684 | 0.897621 | 2.59087 | 3.1248 | 1.09009 | 3.27933 | 2.73522 | 1.11868 | 2.41366 | 3.36125 | 2.04809 | 3.24755 | 4.19625 | 1.57475294 |
| 69 | Hecw2 | NM 001001883 | 0.932703 | 2.88829 | 3.302 | 1.03049 | 2.67371 | 2.70144 | 1.1434 | 2.6016 | 2.24773 | 1.32413 | 1.82133 | 2.4426 | 1.5589107 |
| 70 | Tmem17 | NM 153596 | 6.05101 | 18.1505 | 16.2891 | 5.82476 | 19.2483 | 15.2593 | 7.33446 | 17.1858 | 17.0652 | 9.6624 | 16.747 | 19.618 | 1.54650951 |
| 71 | Dtx3l | NM 001013371 | 0.604163 | 1.78737 | 1.28699 | 0.406379 | 2.30853 | 1.29487 | 0.51412 | 1.25121 | 0.987045 | 0.376359 | 1.1878 | 0.776582 | 1.54554295 |
| 72 | Nefl | NM 010910 | 5.32485 | 13.0672 | 15.2899 | 4.41259 | 11.0256 | 12.7575 | 7.16525 | 9.89302 | 11.6036 | 4.97054 | 9.0766 | 11.6094 | 1.54444081 |
| 73 | Clca4a | NM 207208 | 1.27722 | 9.18266 | 15.9424 | 2.1454 | 9.28571 | 15.0529 | 1.28581 | 6.0026 | 14.3515 | 2.67061 | 7.47146 | 16.8135 | 1.54269493 |
| 74 | Casq1 | NM 009813 | 0.943247 | 2.04231 | 1.85353 | 1.06271 | 2.2597 | 1.79146 | 1.25319 | 1.708 | 1.66092 | 1.07157 | 1.52632 | 2.214 | 1.53967948 |
| 75 | Psmb10 | NM 013640 | 19.2694 | 45.9379 | 57.8198 | 17.5171 | 41.5821 | 60.2537 | 20.1414 | 35.1539 | 54.4991 | 26.6352 | 35.9025 | 54.7763 | 1.5380979 |
| 76 | Hspb2 | NM 024441 | 2.87641 | 6.25495 | 6.01116 | 2.91805 | 6.76297 | 5.12673 | 6.76765 | 10.4806 | 9.80827 | 5.9015 | 8.20076 | 7.37762 | 1.52887596 |
| 77 | Plekha2 | NM 031257 | 3.25944 | 14.2574 | 11.2879 | 3.45934 | 14.8046 | 12.0282 | 5.15204 | 14.4653 | 14.4432 | 4.76934 | 13.9135 | 13.8979 | 1.51158847 |
| 78 | Fgf17 | NM 008004 | 0.605182 | 1.37228 | 2.0613 | 0.694227 | 1.5137 | 2.60155 | 1.25169 | 1.73087 | 2.39176 | 0.957426 | 1.4934 | 1.91392 | 1.51155751 |
| 79 | Nfkbid | NM 172142 | 0.454897 | 1.47357 | 1.40469 | 0.372735 | 1.13991 | 1.22821 | 0.616037 | 1.08621 | 0.686576 | 0.393739 | 0.946408 | 2.3121 | 1.51134734 |
| 80 | Junb | NM 008416 | 3.22879 | 17.445 | 11.9866 | 2.99414 | 17.5976 | 11.1988 | 5.04853 | 18.8297 | 11.6886 | 3.88525 | 14.5243 | 11.4647 | 1.51047316 |
| 81 | Gng11 | NM 025331 | 1.94506 | 6.73828 | 10.2348 | 2.93958 | 6.10761 | 10.668 | 4.13468 | 7.85083 | 11.2135 | 3.44817 | 6.18991 | 9.29111 | 1.50031472 |
| 82 | Cnpy1 | NM 001310512 | 0.437632 | 5.31618 | 2.32672 | 0.463241 | 4.59061 | 2.06516 | 0.318361 | 2.31066 | 1.23152 | 0.276688 | 2.07829 | 1.17504 | 1.4934608 |
| 83 | Spats2l | NM 144882 | 0.687548 | 1.77208 | 2.78445 | 0.462081 | 2.05757 | 3.37788 | 1.08619 | 1.88307 | 3.92726 | 0.74976 | 2.24121 | 2.91176 | 1.48854507 |
| 84 | Apobec2 | NM 009694 | 0.819824 | 3.25858 | 2.7498 | 0.926553 | 4.3252 | 3.48863 | 1.60793 | 3.59117 | 4.77877 | 1.17044 | 4.21998 | 3.77177 | 1.48021383 |
| 85 | Lypd1 | NM 001311089 | 0.719635 | 2.54161 | 2.23902 | 0.659855 | 1.75322 | 1.9049 | 0.590639 | 1.39378 | 1.32751 | 0.761305 | 1.39322 | 1.93044 | 1.47709795 |
| 86 | Smad6 | NM 008542 | 0.12521 | 1.15934 | 0.356763 | 0.235968 | 1.03715 | 0.473288 | 0.431615 | 1.50685 | 0.293967 | 0.202302 | 1.17007 | 0.352672 | 1.47218436 |
| 87 | Kirrel2 | NM 172898 | 0.0502031 | 1.1621 | 0.715224 | 0.0719873 | 1.10077 | 0.962783 | 0.0973447 | 1.02094 | 0.928201 | 0.0633699 | 0.990071 | 1.09775 | 1.47211182 |
| 88 | Miat | NR 033657 | 0.509769 | 2.08685 | 1.12578 | 0.633803 | 1.80966 | 1.10549 | 0.564723 | 1.65912 | 0.948929 | 0.811451 | 1.44687 | 1.19854 | 1.47192456 |
| 89 | Zfp423 | NM 033327 | 0.92075 | 3.50324 | 2.01016 | 0.82443 | 3.50677 | 2.07697 | 1.04092 | 2.60219 | 1.91429 | 0.8759 | 2.60739 | 1.92243 | 1.47138369 |
| 90 | Coch | NM 001198835 | 1.93927 | 10.3667 | 3.3595 | 2.45669 | 9.69781 | 3.90065 | 4.85979 | 13.7762 | 6.60489 | 4.01969 | 14.0129 | 4.63385 | 1.47025492 |

| | | | | | | | | | | | | | | |
|---|---|---|---|---|---|---|---|---|---|---|---|---|---|---|
| 91 | Chst15 | NM 029935 | 0.501998 | 6.12574 | 2.28551 | 0.347602 | 6.67653 | 2.51537 | 0.402149 | 4.0969 | 2.12478 | 0.392689 | 4.39335 | 2.23256 | 1.46945308 |
| 92 | Pitx2 | NM 001287048 | 0.562444 | 59.3012 | 37.9465 | 0.378959 | 62.4512 | 34.8986 | 0.744296 | 48.7914 | 30.8 | 0.433422 | 51.4238 | 40.2304 | 1.4670569 |
| 93 | Cntfr | NM 001136056 | 1.67262 | 3.8644 | 2.31809 | 1.47895 | 3.67025 | 2.36985 | 2.10041 | 3.0811 | 2.10078 | 1.75557 | 3.18522 | 2.06803 | 1.46043103 |
| 94 | Ppp1r36 | NM 001163103 | 0.294637 | 2.58131 | 1.28954 | 0.351158 | 2.08818 | 1.19114 | 0.184665 | 1.13419 | 0.723193 | 0.243433 | 0.975212 | 0.675031 | 1.44931005 |
| 95 | Nodal | NM 013611 | 7.98458 | 57.5156 | 63.441 | 7.36222 | 56.567 | 61.0901 | 8.72146 | 40.7992 | 42.6646 | 7.69382 | 43.5244 | 51.8535 | 1.44040926 |
| 96 | Tcf15 | NM 009328 | 10.4606 | 50.5186 | 23.0266 | 9.2283 | 44.7183 | 23.2188 | 9.14046 | 32.898 | 22.249 | 9.2383 | 28.8407 | 19.0735 | 1.43954178 |
| 97 | Rhob | NM 007483 | 10.9547 | 26.7477 | 19.5212 | 9.02163 | 26.8441 | 17.952 | 13.5806 | 26.7348 | 20.0638 | 13.9785 | 25.427 | 20.9944 | 1.43023988 |
| 98 | Bambi | NM 026505 | 6.77547 | 15.569 | 14.2037 | 7.18384 | 15.1024 | 12.5992 | 10.5726 | 17.0001 | 15.0749 | 11.3525 | 16.7204 | 17.1357 | 1.42825027 |
| 99 | Parp10, Plec | NM 001163575 | 1.6532 | 4.48167 | 4.67359 | 1.74404 | 3.92871 | 4.93412 | 2.08599 | 3.24964 | 3.59836 | 1.76428 | 3.39308 | 2.86715 | 1.42587868 |
| 100 | Psme2 | NM 011190 | 55.2172 | 150.015 | 222.384 | 55.0561 | 138.089 | 214.899 | 59.5933 | 117.952 | 179.447 | 72.9648 | 123.036 | 212.806 | 1.42543659 |
| 101 | Dkk1 | NM 010051 | 0.15884 | 3.40882 | 1.85047 | 0.223116 | 2.32185 | 1.86682 | 0.293327 | 2.90782 | 3.2764 | 0.229141 | 2.86402 | 2.48927 | 1.42186916 |
| 102 | Irf1 | NM 001159396 | 3.98559 | 76.875 | 45.2556 | 3.60987 | 85.2543 | 46.9323 | 4.2965 | 65.2901 | 37.8058 | 3.75993 | 56.3291 | 36.8027 | 1.42176076 |
| 103 | Pik3ip1 | NM 178149 | 1.41147 | 6.55085 | 3.83189 | 1.30674 | 7.15491 | 4.06423 | 1.86231 | 5.48582 | 3.17387 | 1.27394 | 5.33024 | 3.05268 | 1.41891659 |
| 104 | Cx3cl1 | NM 009142 | 0.380528 | 2.13662 | 1.01481 | 0.187079 | 2.48983 | 1.11203 | 0.356042 | 2.36245 | 1.19971 | 0.285453 | 1.91324 | 0.950934 | 1.41881602 |
| 105 | Panct2 | NR 131964 | 0.279958 | 2.86689 | 1.46837 | 0.336136 | 2.97049 | 1.42745 | 0.500375 | 2.98225 | 1.59982 | 0.423832 | 3.18173 | 1.86828 | 1.4166092 |
| 106 | Gm14005 | NR 028590 | 2.96627 | 8.51912 | 11.0191 | 2.64617 | 9.64296 | 8.37072 | 5.39975 | 11.8201 | 13.2295 | 4.95571 | 12.1287 | 13.2975 | 1.40541752 |
| 107 | Dact3 | NM 001081655 | 1.73455 | 5.64871 | 3.481 | 1.79422 | 5.3467 | 3.62428 | 2.02488 | 4.33175 | 2.67159 | 1.88823 | 4.34453 | 3.19143 | 1.40459189 |
| 108 | Trh | NM 009426 | 13.0533 | 73.7802 | 97.784 | 10.9787 | 65.6667 | 87.0471 | 17.1416 | 70.0855 | 100.291 | 13.6232 | 57.4719 | 92.1473 | 1.40039593 |
| 109 | Nxn | NM 008750 | 19.6277 | 54.6674 | 49.9209 | 17.845 | 52.2622 | 53.1287 | 15.4158 | 30.882 | 40.3562 | 13.2334 | 27.7859 | 34.9688 | 1.39263041 |
| 110 | Radil | NM 178702 | 0.526607 | 2.69195 | 2.28767 | 0.292919 | 2.64475 | 2.61923 | 0.484543 | 1.79989 | 2.53374 | 0.324418 | 2.1166 | 2.47389 | 1.38108644 |
| 111 | Prkd2 | NM 178900 | 0.725773 | 1.65623 | 1.09259 | 0.865614 | 1.84072 | 1.35492 | 0.967959 | 1.57962 | 1.16732 | 0.816239 | 1.28181 | 1.19825 | 1.37667347 |
| 112 | Fgf3 | NM 008007 | 1.42167 | 3.94009 | 3.0963 | 1.20372 | 3.69446 | 2.58156 | 1.34769 | 2.77233 | 1.72453 | 1.53134 | 3.36447 | 1.65092 | 1.37292334 |
| 113 | Abcg2 | NM 011920 | 29.7662 | 88.0869 | 107.092 | 27.6785 | 79.3663 | 98.1564 | 29.982 | 62.7143 | 95.1259 | 32.9048 | 70.9391 | 102.855 | 1.37176269 |

| | | | | | | | | | | | | | | | |
|---|---|---|---|---|---|---|---|---|---|---|---|---|---|---|---|
| 114 | Cdh6 | NM 007666 | 0.590997 | 1.25636 | 1.21482 | 0.477882 | 1.00788 | 1.14118 | 0.797477 | 1.15248 | 1.05892 | 0.526126 | 0.868245 | 0.818484 | 1.36811484 |
| 115 | Rab19 | NM 011226 | 1.11144 | 3.42334 | 4.78304 | 1.51176 | 3.37928 | 6.32653 | 1.36011 | 1.80821 | 3.41837 | 0.830541 | 2.13121 | 4.02028 | 1.36450017 |
| 116 | Gm15417 | NR 040403 | 2.45273 | 5.74517 | 5.91003 | 2.39882 | 5.90047 | 4.68175 | 2.88734 | 5.12216 | 4.78992 | 3.17419 | 5.56203 | 4.40929 | 1.3618045 |
| 117 | Pycr2 | NM 133705 | 42.1748 | 196.726 | 158.516 | 38.38 | 192.002 | 174.435 | 41.9345 | 153.716 | 158.964 | 44.6803 | 156.345 | 174.138 | 1.34925986 |
| 118 | Mfap4 | NM 029568 | 2.52537 | 7.81538 | 6.11613 | 2.33416 | 6.85444 | 6.78938 | 2.39258 | 4.67624 | 5.06213 | 1.24862 | 3.15523 | 2.99781 | 1.34584201 |
| 119 | Sfn | NM 018754 | 3.31141 | 13.2777 | 8.54386 | 3.50746 | 13.4851 | 10.0268 | 4.22259 | 13.7399 | 10.8401 | 3.83327 | 9.89995 | 9.41889 | 1.34572373 |
| 120 | Dok2 | NM 010071 | 3.02323 | 10.1218 | 10.7898 | 3.43606 | 11.95 | 10.7023 | 3.19902 | 8.49958 | 8.08047 | 3.12967 | 7.5878 | 11.182 | 1.34329555 |
| 121 | Gm15867 | NR 131748 | 0.189811 | 1.31513 | 1.11897 | 0.492506 | 1.079 | 0.66824 | 0.29989 | 1.1633 | 1.5226 | 0.44724 | 1.30536 | 1.50072 | 1.34153376 |
| 122 | Pdlim3 | NM 016798 | 3.43725 | 8.63893 | 10.9483 | 3.11096 | 8.73527 | 11.9387 | 4.02701 | 7.3099 | 9.99517 | 3.35756 | 7.22517 | 10.1245 | 1.34132978 |
| 123 | Mpc1 | NM 018819 | 15.109 | 40.0859 | 26.2412 | 17.5489 | 36.3881 | 24.4328 | 18.5447 | 33.7859 | 29.0234 | 22.3466 | 38.2167 | 31.3728 | 1.33821721 |
| 124 | Stx3 | NM 001025307 | 18.5157 | 48.4691 | 53.7859 | 18.057 | 44.5292 | 56.1264 | 16.2585 | 31.6431 | 46.1631 | 20.3795 | 37.9778 | 53.7422 | 1.3343989 |
| 125 | Mir9-3hg | NR 040314 | 1.25942 | 2.57227 | 0.913724 | 1.00436 | 2.53972 | 0.921418 | 1.19223 | 1.46157 | 1.548 | 0.978653 | 2.15383 | 0.828313 | 1.33396208 |
| 126 | Krt42 | NM 212483 | 0.854938 | 3.39733 | 3.16235 | 1.17441 | 3.35707 | 3.32669 | 0.888462 | 2.21356 | 2.45 | 0.863331 | 2.28876 | 2.31115 | 1.32858477 |
| 127 | Lmo2 | NM 001142337 | 0.401311 | 2.10354 | 2.0495 | 0.641758 | 2.15326 | 2.15258 | 0.468057 | 1.18021 | 1.55861 | 0.378002 | 1.49293 | 1.50761 | 1.32852288 |
| 128 | Six2 | NM 011380 | 0.819342 | 2.82645 | 1.29178 | 1.06168 | 2.58104 | 1.54642 | 1.19872 | 2.54793 | 1.81181 | 1.07753 | 2.49749 | 1.15814 | 1.32349918 |
| 129 | Zfp870 | NM 207245 | 0.848176 | 2.02011 | 1.38892 | 0.709461 | 1.85974 | 1.51267 | 0.670034 | 1.2208 | 0.891958 | 0.678064 | 1.32959 | 1.20499 | 1.32255861 |
| 130 | Syt13 | NM 030725 | 2.65262 | 10.2799 | 8.87961 | 2.13539 | 9.43296 | 8.96102 | 2.53464 | 7.83044 | 8.50044 | 2.02075 | 6.51762 | 7.79577 | 1.31325247 |
| 131 | Ctgf | NM 010217 | 6.65183 | 22.3851 | 21.0103 | 5.93515 | 22.3855 | 21.4958 | 9.42559 | 25.5972 | 28.8122 | 7.59138 | 20.8036 | 24.6846 | 1.30805664 |
| 132 | Aldh1b1 | NM 028270 | 1.76961 | 3.58754 | 1.75365 | 1.28691 | 3.91934 | 1.36547 | 1.38335 | 2.34856 | 1.23596 | 0.974442 | 2.12703 | 1.1898 | 1.30724916 |
| 133 | 1110037F02 Rik | NM 001081183 | 0.559263 | 1.8494 | 1.02048 | 0.752842 | 1.67394 | 1.24353 | 0.895078 | 1.82508 | 1.40683 | 1.04883 | 2.32708 | 1.65816 | 1.29888736 |
| 134 | Avpi1 | NM 027106 | 44.0682 | 106.622 | 88.3415 | 40.475 | 101.647 | 76.8091 | 50.0119 | 107.801 | 82.5406 | 67.1997 | 110.606 | 109.774 | 1.29709629 |
| 135 | Nedd9 | NM 001111324 | 1.03481 | 3.44705 | 1.81119 | 1.21683 | 3.04797 | 2.27713 | 1.36056 | 3.25313 | 2.41069 | 1.57683 | 3.33766 | 2.403 | 1.29465643 |
| 136 | Steap1 | NM 027399 | 6.38516 | 15.1233 | 14.5238 | 6.73697 | 15.0303 | 16.5313 | 7.81425 | 14.0352 | 15.6756 | 7.30167 | 12.8445 | 16.5109 | 1.29373811 |

| | | | | | | | | | | | | | | |
|---|---|---|---|---|---|---|---|---|---|---|---|---|---|---|
| 137 | Rps6kl1 | NM 146244 | 1.86834 | 7.10777 | 6.79042 | 1.61217 | 6.65442 | 8.62102 | 1.73834 | 5.74831 | 6.1139 | 1.51936 | 4.29411 | 5.29432 | 1.29331226 |
| 138 | Igsf11 | NM 170599 | 0.506952 | 1.04041 | 2.38185 | 0.200394 | 1.28149 | 2.28455 | 0.608473 | 1.50506 | 1.67861 | 0.269919 | 1.10449 | 1.05398 | 1.28660746 |
| 139 | Mup6 | NM 001081285 | 7.031 | 23.0696 | 36.7066 | 8.94901 | 23.2067 | 37.0421 | 6.91061 | 18.414 | 34.018 | 12.227 | 23.5425 | 44.5393 | 1.27979878 |
| 140 | Mb21d2 | NM 177718 | 4.48131 | 10.8491 | 9.22452 | 4.80622 | 10.0486 | 9.44137 | 5.35318 | 8.84016 | 8.88576 | 5.54686 | 10.4034 | 10.4978 | 1.27921796 |
| 141 | Homer3 | NM 001146153 | 8.51853 | 20.0667 | 16.8077 | 7.0828 | 16.046 | 14.7205 | 9.82977 | 16.8344 | 13.9449 | 7.35711 | 14.0646 | 14.631 | 1.27504545 |
| 142 | Lgr4 | NM 172671 | 7.21139 | 24.0115 | 17.3459 | 6.75983 | 24.4141 | 18.2935 | 8.28253 | 21.5198 | 19.4599 | 7.00862 | 20.0323 | 15.6246 | 1.27212807 |
| 143 | Kazn | NM 001109685 | 0.900672 | 2.15805 | 1.46766 | 0.668391 | 2.3171 | 1.72682 | 0.866703 | 2.05637 | 1.36049 | 0.911429 | 2.05209 | 1.20958 | 1.2678516 |
| 144 | Irgm1 | NM 008326 | 0.342823 | 5.80374 | 3.62634 | 0.281093 | 5.45 | 4.32928 | 0.251043 | 4.3469 | 3.63559 | 0.363448 | 4.12411 | 3.20467 | 1.26708453 |
| 145 | Pcolce2 | NM 029620 | 7.5946 | 16.0068 | 13.6824 | 7.00215 | 14.1664 | 14.0017 | 6.84934 | 11.6723 | 12.5197 | 6.39325 | 9.97542 | 10.4127 | 1.26538916 |
| 146 | Mdfi | NM 010783 | 8.14743 | 25.8088 | 13.0969 | 7.43034 | 24.8124 | 14.45 | 7.86349 | 19.2484 | 13.6456 | 7.56159 | 20.486 | 15.729 | 1.26178228 |
| 147 | Relt | NM 177073 | 1.69514 | 5.47778 | 3.41366 | 1.45988 | 5.01333 | 3.27271 | 1.74249 | 4.82558 | 2.6628 | 1.67762 | 4.25333 | 2.88183 | 1.25653448 |
| 148 | Notch3 | NM 008716 | 1.05644 | 4.4421 | 4.4649 | 0.849112 | 4.86462 | 4.74285 | 0.96255 | 3.79673 | 3.76307 | 0.785859 | 3.127 | 3.46101 | 1.25371437 |
| 149 | Bcar3 | NM 013867 | 2.74729 | 24.9068 | 15.656 | 2.55827 | 25.175 | 18.275 | 2.74169 | 22.3539 | 14.3865 | 2.71765 | 18.8377 | 13.7737 | 1.25334165 |
| 150 | Prrx2 | NM 009116 | 3.11648 | 8.96375 | 6.36792 | 2.4858 | 7.95539 | 5.24313 | 5.98943 | 12.2622 | 9.55078 | 4.59528 | 12.9533 | 8.91296 | 1.24874817 |
| 151 | Rnd3 | NM 028810 | 18.7578 | 42.4265 | 38.8494 | 19.5174 | 42.9093 | 41.7987 | 23.4656 | 41.0781 | 46.7169 | 25.4283 | 46.8712 | 49.6464 | 1.24110311 |
| 152 | Btg2 | NM 007570 | 9.01833 | 20.6619 | 9.88989 | 7.74535 | 19.6681 | 9.01787 | 8.38834 | 16.3835 | 8.92215 | 7.44286 | 14.5414 | 7.72612 | 1.23639854 |
| 153 | Nxph3 | NM 130858 | 0.495301 | 1.68042 | 0.420285 | 0.295158 | 1.57966 | 0.525623 | 0.388039 | 1.42998 | 0.581434 | 0.38649 | 1.31151 | 0.43388 | 1.23537457 |
| 154 | Tppp3 | NM 026481 | 3.53584 | 15.4857 | 14.9596 | 4.06072 | 17.1942 | 12.671 | 4.59389 | 17.4111 | 13.067 | 6.27818 | 20.0489 | 14.7155 | 1.23346959 |
| 155 | C130071C03 Rik | NR 015561 | 0.296566 | 1.20486 | 0.509922 | 0.328053 | 1.19937 | 0.741845 | 0.357807 | 1.20621 | 0.57443 | 0.389321 | 1.15174 | 0.615503 | 1.21949469 |
| 156 | Hey1 | NM 010423 | 1.07244 | 2.61389 | 1.83251 | 0.776854 | 2.68089 | 2.15994 | 1.52696 | 2.21687 | 1.73331 | 0.878927 | 2.98031 | 1.86632 | 1.21591756 |
| 157 | Unc5b | NM 029770 | 2.1757 | 10.5298 | 5.65503 | 2.02831 | 10.8932 | 5.75091 | 2.42983 | 10.1058 | 5.32311 | 2.22508 | 9.43697 | 5.06842 | 1.21547845 |
| 158 | Igtp,Irgm2 | NM 018738 | 2.18732 | 4.48413 | 3.65279 | 2.22257 | 5.15513 | 4.62806 | 1.92617 | 3.60189 | 3.63713 | 1.63993 | 2.83045 | 2.9548 | 1.21512338 |
| 159 | Tmem119 | NM 146162 | 0.483875 | 2.66683 | 0.950839 | 0.46256 | 2.92354 | 0.96831 | 0.430897 | 2.22865 | 0.852032 | 0.407188 | 1.85959 | 1.00425 | 1.21488033 |

| | | | | | | | | | | | | | | | |
|---|---|---|---|---|---|---|---|---|---|---|---|---|---|---|---|
| 160 | Platr8 | NR 033473 | 1.8247 | 4.66493 | 4.74568 | 1.84692 | 5.64832 | 4.41522 | 3.2375 | 6.81751 | 4.03203 | 2.27617 | 5.73162 | 2.86414 | 1.21429795 |
| 161 | Trim34b | NM 001243916 | 0.56013 | 1.71996 | 0.997494 | 0.803183 | 1.77401 | 1.55682 | 0.633559 | 1.36535 | 1.35617 | 0.618656 | 1.35957 | 1.73857 | 1.21290448 |
| 162 | Zfp946 | NM 198003 | 5.37696 | 15.823 | 16.2514 | 6.40525 | 15.0257 | 16.4691 | 4.82762 | 11.4119 | 15.3559 | 8.26133 | 16.572 | 21.561 | 1.21024357 |
| 163 | Nkx6-2 | NM 183248 | 2.84058 | 6.38847 | 6.55786 | 1.98482 | 6.30053 | 6.18044 | 2.58314 | 6.03933 | 5.03253 | 2.46371 | 5.3297 | 5.01189 | 1.2048532 |
| 164 | Pkdcc | NM 134117 | 4.63022 | 12.8644 | 12.7435 | 3.76016 | 13.5097 | 13.1365 | 5.68345 | 14.1772 | 16.1611 | 5.49427 | 15.4907 | 15.5002 | 1.19897057 |
| 165 | Pde4b | NM 001177983 | 0.808071 | 3.26848 | 3.48733 | 0.750579 | 3.01661 | 3.02964 | 0.666832 | 1.7842 | 3.36221 | 0.604623 | 2.45521 | 3.77366 | 1.19706056 |
| 166 | Lrrc2 | NM 028838 | 1.1401 | 2.52486 | 2.21211 | 1.0219 | 2.85821 | 2.20181 | 0.877736 | 2.09183 | 1.98433 | 1.00735 | 1.82577 | 1.57346 | 1.19446108 |
| 167 | 1700057H15 Rik | NR 040774 | 0.748667 | 2.12205 | 1.10338 | 0.920169 | 2.00634 | 0.62425 | 0.806488 | 1.45994 | 1.53801 | 0.756016 | 1.81719 | 1.10986 | 1.19007598 |
| 168 | Cacna1g | NM 001177890 | 0.850654 | 7.39594 | 3.92038 | 1.11049 | 8.5047 | 4.46207 | 0.715599 | 6.18275 | 2.51363 | 1.16677 | 5.9615 | 3.01782 | 1.18935838 |
| 169 | Msx2 | NM 013601 | 0.678747 | 1.77958 | 1.28376 | 0.472732 | 1.53785 | 0.98098 | 1.16987 | 2.70391 | 1.85252 | 0.97671 | 2.57419 | 1.81118 | 1.18761561 |
| 170 | Dok5 | NM 029761 | 0.521039 | 1.70248 | 2.23971 | 0.462509 | 1.22707 | 2.05157 | 0.374187 | 0.798329 | 2.34465 | 0.350769 | 1.00121 | 1.57701 | 1.18699763 |
| 171 | Crlf1 | NM 018827 | 2.78933 | 10.819 | 10.0689 | 2.27731 | 10.103 | 9.06792 | 2.65524 | 8.47013 | 8.84173 | 1.90345 | 7.29694 | 8.23668 | 1.18389433 |
| 172 | Kif26a | NM 001097621 | 0.808401 | 1.91867 | 0.794277 | 0.873879 | 1.95049 | 0.747158 | 0.74286 | 1.51402 | 0.705427 | 0.899938 | 1.67259 | 0.840161 | 1.18188595 |
| 173 | Trim6 | NM 001013616 | 2.12058 | 4.9178 | 2.65176 | 2.36941 | 6.38741 | 3.00054 | 1.99362 | 4.94081 | 5.205 | 3.69878 | 6.55092 | 2.85787 | 1.18013064 |
| 174 | Kbtbd3 | NM 026962 | 2.47347 | 6.18086 | 5.97665 | 2.72101 | 7.08988 | 5.1245 | 2.94012 | 6.95768 | 6.32599 | 4.31941 | 8.46464 | 8.43794 | 1.17991381 |
| 175 | Plekhh1 | NM 181073 | 0.929214 | 1.90297 | 1.10144 | 0.861404 | 1.94274 | 0.988051 | 0.874851 | 1.47057 | 1.08568 | 0.660356 | 1.30004 | 1.35073 | 1.17909192 |
| 176 | Pde1b | NM 008800 | 1.00492 | 3.29703 | 2.56062 | 0.542911 | 3.28227 | 2.30811 | 0.732791 | 2.64568 | 2.19083 | 0.591029 | 2.5598 | 1.99724 | 1.17440889 |
| 177 | Nespas | NR 002846 | 1.03137 | 2.08798 | 1.71593 | 0.779441 | 1.65037 | 1.36589 | 1.05126 | 1.53759 | 1.27925 | 0.815073 | 1.6866 | 1.36363 | 1.1727041 |
| 178 | Pth1r | NM 001083936 | 0.843698 | 2.44705 | 0.929381 | 0.614662 | 2.59598 | 1.15562 | 1.1506 | 2.73098 | 1.57001 | 0.68287 | 2.54962 | 1.29268 | 1.16645954 |
| 179 | Tnfaip6 | NM 009398 | 0.909185 | 3.77565 | 3.59397 | 0.987522 | 3.72946 | 3.1264 | 1.89503 | 7.18609 | 5.88252 | 1.94298 | 5.85061 | 5.48532 | 1.16553099 |
| 180 | Runx1 | NM 009821 | 1.09983 | 2.5023 | 1.8065 | 0.399267 | 3.42325 | 3.65475 | 1.09937 | 4.00112 | 2.0737 | 0.576252 | 3.28511 | 2.16615 | 1.16152802 |
| 181 | Ubr7 | NM 025666 | 51.9925 | 144.464 | 127.229 | 49.0124 | 147.54 | 126.657 | 49.9152 | 119.86 | 123.974 | 51.047 | 131.937 | 130.583 | 1.16103895 |
| 182 | Efna3 | NM 010108 | 1.30428 | 3.80203 | 1.34822 | 0.871107 | 4.85718 | 1.93922 | 1.2044 | 4.54822 | 1.60402 | 1.27009 | 4.50291 | 1.91101 | 1.15969386 |

| | | | | | | | | | | | | | | | |
|---|---|---|---|---|---|---|---|---|---|---|---|---|---|---|---|
| 183 | Papolb | NM 019943 | 0.312271 | 1.76395 | 1.04448 | 0.391971 | 1.89708 | 0.775589 | 0.326368 | 1.33264 | 0.935935 | 0.261662 | 1.2982 | 0.709162 | 1.15965554 |
| 184 | Fermt1 | NM 198029 | 0.385989 | 1.26216 | 1.27098 | 0.362832 | 1.1945 | 1.377 | 0.276531 | 0.765179 | 0.949246 | 0.2333 | 0.677595 | 0.875258 | 1.15703854 |
| 185 | Fat1 | NM 001081286 | 4.73375 | 13.5995 | 9.99964 | 5.25566 | 13.522 | 9.99971 | 6.34218 | 14.9884 | 11.6141 | 4.98684 | 11.7158 | 8.67329 | 1.15555944 |
| 186 | Fam25c | NM 183278 | 19.6348 | 49.2816 | 44.7768 | 15.7274 | 48.1506 | 42.9726 | 19.8784 | 52.3374 | 52.1702 | 24.2645 | 53.1242 | 52.7759 | 1.15536864 |
| 187 | Arl4a | NM 007487 | 93.2095 | 241.082 | 211.314 | 95.0883 | 228.516 | 197.921 | 98.1099 | 230.455 | 197.801 | 118.51 | 234.529 | 227.353 | 1.15289598 |
| 188 | Lama5 | NM 001081171 | 1.65631 | 7.11897 | 4.41595 | 1.93912 | 7.6573 | 4.35696 | 1.86044 | 6.88209 | 4.07982 | 1.68391 | 5.85662 | 3.94111 | 1.14905354 |
| 189 | Yaf2 | NR 028315 | 9.90559 | 28.0411 | 23.4499 | 10.9475 | 26.3159 | 22.704 | 12.1679 | 27.0425 | 24.9587 | 13.2182 | 30.9 | 30.6974 | 1.14791928 |
| 190 | S100a10 | NM 009112 | 128.409 | 264.703 | 281.956 | 132.73 | 272.495 | 256.456 | 201.629 | 407.468 | 367.933 | 243.364 | 383.281 | 424.685 | 1.14422314 |
| 191 | Slc7a7 | NM 011405 | 36.6397 | 84.2148 | 85.0234 | 38.1061 | 77.194 | 92.4098 | 30.9626 | 58.7557 | 73.9918 | 30.8786 | 58.7152 | 67.9434 | 1.13821698 |
| 192 | Nfic | NM 008688 | 1.74575 | 5.20568 | 1.58376 | 1.68893 | 4.04038 | 1.97687 | 1.93094 | 3.22062 | 1.29973 | 1.21491 | 3.75783 | 1.61377 | 1.12879492 |
| 193 | Eomes | NM 010136 | 1.18592 | 5.48914 | 4.00385 | 1.20438 | 4.99856 | 4.03658 | 1.11013 | 4.11732 | 3.33213 | 0.900897 | 3.67741 | 2.99994 | 1.12682956 |
| 194 | Pdlim4 | NM 019417 | 0.456595 | 1.12174 | 1.34585 | 0.395859 | 1.58912 | 0.979033 | 0.562164 | 1.92615 | 1.07201 | 0.562471 | 1.3062 | 1.15567 | 1.12569095 |
| 195 | Nectin4 | NM 027893 | 0.420467 | 1.25731 | 0.54687 | 0.429637 | 1.3799 | 0.545299 | 0.290581 | 0.902099 | 0.455186 | 0.356589 | 0.858398 | 0.373918 | 1.12524912 |
| 196 | Mrc2 | NM 008626 | 0.605485 | 1.637 | 1.12574 | 0.561003 | 1.94716 | 1.36184 | 0.626158 | 1.62967 | 1.21926 | 0.573803 | 1.6675 | 1.15998 | 1.12085851 |
| 197 | Dok1 | NM 001291799 | 5.20029 | 12.0674 | 8.48445 | 4.431 | 11.8727 | 8.04994 | 5.37806 | 11.8792 | 9.29812 | 5.46802 | 12.3887 | 10.0043 | 1.11744271 |
| 198 | Sertad4 | NM 001177794 | 2.94227 | 10.5338 | 9.65874 | 2.9672 | 11.0663 | 10.0211 | 2.48472 | 8.31168 | 8.19009 | 2.5538 | 8.16864 | 7.64403 | 1.11705309 |
| 199 | Irak2 | NM 172161 | 1.30834 | 5.4189 | 4.28029 | 1.31847 | 5.6365 | 4.38039 | 2.08723 | 8.78521 | 6.31473 | 2.46701 | 8.24584 | 6.75847 | 1.11459662 |
| 200 | Card10 | NM 130859 | 0.988089 | 2.26158 | 1.66134 | 0.695593 | 2.25128 | 2.0191 | 1.15735 | 2.86117 | 1.80523 | 0.857256 | 2.13654 | 1.50605 | 1.11297433 |
| 201 | Endod1 | NM 028013 | 0.600349 | 1.88345 | 1.20997 | 0.636969 | 1.80013 | 1.15519 | 0.729628 | 2.01552 | 1.5631 | 0.551837 | 1.44785 | 1.18665 | 1.10717624 |
| 202 | Hes1 | NM 008235 | 4.72298 | 21.2382 | 16.8501 | 4.09454 | 23.1689 | 18.1924 | 4.53121 | 18.8629 | 15.269 | 3.88252 | 19.899 | 15.4056 | 1.09335572 |
| 203 | Msx1 | NM 010835 | 2.34499 | 17.8448 | 5.96542 | 2.0555 | 17.821 | 6.40648 | 2.94516 | 18.6155 | 7.76754 | 2.00088 | 17.1638 | 7.57246 | 1.09268043 |
| 204 | Kras | NM 021284 | 10.266 | 24.2726 | 16.9592 | 10.0246 | 23.8408 | 17.582 | 10.3242 | 21.181 | 18.1208 | 9.3519 | 21.5232 | 18.3834 | 1.08948428 |
| 205 | Thbs1 | NM 001313914 | 1.48261 | 5.67241 | 4.47535 | 1.21021 | 5.25187 | 4.5438 | 1.70035 | 6.26291 | 6.23382 | 1.29223 | 4.9396 | 5.86664 | 1.08789841 |

| | | | | | | | | | | | | | | |
|---|---|---|---|---|---|---|---|---|---|---|---|---|---|---|
| 206 | Cpt1a | NM 013495 | 1.4929 | 5.33466 | 2.26606 | 1.02537 | 6.21886 | 3.32895 | 0.854255 | 3.7169 | 2.55693 | 0.872327 | 3.96314 | 1.80569 | 1.08366336 |
| 207 | Lax1 | NM 001159649 | 0.758729 | 2.71159 | 1.653 | 0.810901 | 2.89324 | 1.50367 | 0.824434 | 3.67593 | 1.66517 | 1.79885 | 3.96348 | 1.56491 | 1.07200734 |
| 208 | Pxdc1 | NM 025831 | 2.64308 | 11.8682 | 10.1395 | 2.93497 | 10.8281 | 7.72318 | 2.63265 | 7.99053 | 10.8901 | 1.68686 | 8.02567 | 9.13248 | 1.04962294 |
| 209 | Dmtn | NM 001252663 | 0.980323 | 3.2199 | 1.90834 | 1.105 | 3.24825 | 1.71437 | 1.3803 | 3.48166 | 1.78852 | 0.672975 | 2.32274 | 1.15498 | 1.04189565 |
| 210 | Ap3b2 | NM 021492 | 0.540465 | 1.18541 | 1.38445 | 0.569377 | 1.31671 | 1.50216 | 0.615475 | 1.09695 | 1.37087 | 0.331361 | 0.843324 | 1.45935 | 1.04126053 |
| 211 | Il17rd | NM 134437 | 1.55254 | 4.00021 | 3.83195 | 1.39197 | 4.39635 | 4.23173 | 1.32503 | 3.79853 | 3.77272 | 1.24968 | 3.30872 | 3.5135 | 1.0399898 |
| 212 | Sema3f | NM 011349 | 0.860605 | 1.72224 | 1.33961 | 0.83118 | 1.85504 | 1.38075 | 0.859589 | 1.61013 | 1.14965 | 0.742596 | 1.63475 | 1.37961 | 1.03889363 |
| 213 | Rnf11 | NM 013876 | 8.80517 | 17.7477 | 16.6784 | 8.67096 | 17.5923 | 16.1454 | 8.73852 | 18.0196 | 16.8214 | 9.89851 | 18.2591 | 19.4904 | 1.03526133 |
| 214 | Zmiz1 | NM 183208 | 0.939801 | 2.79921 | 1.50914 | 0.845769 | 3.20399 | 1.32714 | 1.00892 | 3.13245 | 1.61883 | 0.889175 | 3.06882 | 1.81268 | 1.03213862 |
| 215 | Tle3 | NM 009389 | 3.6028 | 7.70373 | 3.38907 | 2.92651 | 9.03765 | 3.57082 | 3.48375 | 8.68119 | 3.47532 | 3.06938 | 7.974 | 3.58165 | 1.02684455 |
| 216 | Nav1 | NM 173437 | 1.61618 | 4.1193 | 2.00896 | 1.44097 | 3.03029 | 2.13812 | 1.36666 | 3.19612 | 1.95584 | 1.33305 | 2.94845 | 1.92888 | 1.0222607 |
| 217 | Tbc1d9 | NM 001111304 | 0.707254 | 1.90299 | 0.983774 | 0.540364 | 2.67384 | 0.958107 | 0.700759 | 2.9572 | 1.40076 | 0.675509 | 2.21299 | 1.36243 | 1.01905877 |
| 218 | Cthrc1 | NM 026778 | 9.36431 | 19.4532 | 22.5593 | 8.45994 | 18.3974 | 23.8915 | 6.5181 | 15.0168 | 20.9638 | 8.15298 | 15.4808 | 21.4906 | 1.01174825 |
| 219 | Mmp25 | NM 001033339 | 1.5638 | 5.59246 | 5.11484 | 1.43309 | 5.80831 | 5.62745 | 1.15851 | 3.96663 | 3.88888 | 1.05277 | 4.34509 | 3.98577 | 1.01032909 |
| 220 | Nuak1 | NM 001004363 | 0.671403 | 2.29131 | 2.16668 | 0.621275 | 2.26179 | 1.95617 | 0.86292 | 3.18611 | 2.52755 | 0.695528 | 2.28995 | 2.26521 | 1.00982838 |
| 221 | Ncor2 | NM 001253904 | 2.08287 | 4.31905 | 3.27071 | 2.26282 | 4.64078 | 2.96334 | 2.18693 | 4.46534 | 2.92003 | 2.15984 | 4.45591 | 3.00621 | 1.00477091 |
| 222 | Cxxc5 | NM 133687 | 9.58335 | 19.3478 | 12.5177 | 7.84663 | 19.4038 | 11.955 | 9.67464 | 20.7708 | 11.7691 | 8.49347 | 20.0595 | 11.7734 | 0.99625001 |
| 223 | Cpeb1 | NM 001252525 | 0.557026 | 1.22094 | 1.60302 | 0.638015 | 1.43287 | 1.28663 | 0.657767 | 1.30093 | 2.02608 | 0.516336 | 1.28674 | 1.68102 | 0.99280926 |
| 224 | Snhg9 | NR 027900 | 29.7743 | 61.707 | 50.0149 | 31.4795 | 87.0456 | 45.9153 | 29.7976 | 84.5504 | 56.3743 | 56.2536 | 116.563 | 78.311 | 0.98534607 |
| 225 | Atp1b2 | NM 013415 | 0.599138 | 1.31721 | 0.803758 | 0.679741 | 1.4372 | 0.883855 | 0.804281 | 1.69475 | 0.784481 | 0.605019 | 1.38433 | 0.631982 | 0.98125353 |
| 226 | Whrn | NM 001008793 | 1.28372 | 2.98047 | 2.32644 | 1.26404 | 2.7924 | 2.64564 | 1.14433 | 2.25716 | 1.64309 | 0.91088 | 2.42003 | 1.62937 | 0.97873848 |
| 227 | Ago1 | NM 001317173 | 2.31317 | 6.51648 | 4.79783 | 2.28601 | 6.76925 | 4.89793 | 2.13162 | 5.61368 | 3.83663 | 1.51347 | 4.96327 | 3.71188 | 0.97722948 |
| 228 | Camk2n1 | NM 025451 | 2.08228 | 6.48927 | 1.57176 | 2.1165 | 5.76052 | 1.579 | 1.56176 | 4.61839 | 1.63648 | 1.37553 | 4.20991 | 1.02867 | 0.97015525 |

| | | | | | | | | | | | | | | | |
|---|---|---|---|---|---|---|---|---|---|---|---|---|---|---|---|
| 229 | Tcirg1 | NM 001136091 | 0.932486 | 3.35634 | 1.81851 | 0.78387 | 3.13577 | 1.76071 | 0.715537 | 2.69272 | 1.43954 | 0.696557 | 2.84291 | 1.66607 | 0.96878436 |
| 230 | Epb41l1 | NM 013510 | 0.7644 | 2.60771 | 2.12366 | 1.00025 | 2.02535 | 1.87312 | 0.806007 | 2.38748 | 1.19759 | 1.04246 | 2.77114 | 1.91373 | 0.96724643 |
| 231 | Itga5 | NM 010577 | 2.16227 | 4.83519 | 3.15365 | 1.79625 | 4.83403 | 3.11095 | 2.74032 | 6.51136 | 4.55838 | 1.91906 | 5.24835 | 3.62694 | 0.96406894 |
| 232 | Bhlhe40 | NM 011498 | 0.450563 | 1.72356 | 2.32142 | 0.450225 | 1.59567 | 2.12584 | 0.61558 | 1.91032 | 2.39948 | 0.388403 | 1.77381 | 2.28076 | 0.96079539 |
| 233 | Foxp4 | NM 001110825 | 3.31912 | 6.7766 | 3.71069 | 2.31723 | 7.59518 | 3.88315 | 2.65463 | 7.05373 | 2.72987 | 2.10787 | 6.09111 | 2.66328 | 0.95899327 |
| 234 | Jag1 | NM 013822 | 0.671902 | 1.9967 | 1.16234 | 0.569369 | 1.41028 | 1.04706 | 0.633475 | 1.683 | 1.05166 | 0.504525 | 1.52633 | 0.868361 | 0.95891895 |
| 235 | Rem2 | NM 080726 | 0.544227 | 1.12396 | 1.11403 | 0.436099 | 1.03731 | 1.41016 | 0.53842 | 0.848097 | 1.03257 | 0.411 | 1.26373 | 1.66468 | 0.95568206 |
| 236 | Irf2bpl | NM 145836 | 4.58893 | 10.8037 | 6.74094 | 3.88955 | 10.7546 | 6.55703 | 4.46684 | 11.6521 | 6.35837 | 4.25415 | 11.7775 | 6.64673 | 0.95206361 |
| 237 | Epha2 | NM 010139 | 2.79911 | 14.3277 | 12.7364 | 2.22473 | 14.3456 | 12.3148 | 2.09602 | 12.8853 | 10.1682 | 1.87538 | 11.289 | 10.3294 | 0.95067166 |
| 238 | Agrn | NM 021604 | 3.68072 | 8.89625 | 7.28749 | 4.29034 | 9.73661 | 7.85968 | 4.162 | 10.6033 | 7.36426 | 3.85689 | 9.33441 | 7.61778 | 0.94335085 |
| 239 | Chd9 | NM 177224 | 4.27598 | 9.16509 | 7.80629 | 4.56325 | 9.4187 | 7.86699 | 3.49571 | 8.75585 | 6.74941 | 3.58086 | 7.12465 | 6.19929 | 0.93615003 |
| 240 | Map2k4 | NM 001316367 | 2.3137 | 7.15422 | 6.40649 | 2.92989 | 6.55916 | 5.8465 | 2.71897 | 9.84402 | 7.40742 | 4.03562 | 8.42385 | 9.35439 | 0.93394156 |
| 241 | Ccdc85a | NM 181577 | 1.55763 | 3.85304 | 2.92517 | 1.25362 | 3.00643 | 2.26381 | 1.27813 | 3.41135 | 3.40645 | 1.11298 | 2.87251 | 2.81764 | 0.92798385 |
| 242 | Zfhx3 | NM 007496 | 3.81068 | 9.51877 | 4.84723 | 4.26846 | 10.8046 | 4.3722 | 3.75142 | 10.1833 | 5.30742 | 3.24303 | 8.81635 | 4.4441 | 0.92566105 |
| 243 | Lrrn4 | NM 177303 | 1.41771 | 3.15913 | 2.75083 | 1.37352 | 3.53646 | 2.60709 | 1.42155 | 3.04033 | 2.47021 | 1.17952 | 3.62508 | 2.72294 | 0.92152493 |
| 244 | Gm15706 | NR_045598 | 0.610117 | 1.7151 | 1.95958 | 0.844172 | 1.83425 | 1.90911 | 0.807477 | 1.76873 | 1.66182 | 0.616704 | 2.01825 | 2.11002 | 0.91229481 |
| 245 | Tmem63a | NM 144794 | 0.15667 | 1.85468 | 1.08389 | 0.231073 | 1.89278 | 1.39346 | 0.180021 | 1.55149 | 1.1638 | 0.105471 | 1.48585 | 0.884714 | 0.88211354 |
| 246 | Trim47 | NM 001205081 | 0.643872 | 2.11742 | 1.21474 | 0.506571 | 2.2064 | 1.20282 | 0.531486 | 2.4949 | 0.967659 | 0.494114 | 1.96942 | 0.992393 | 0.88066471 |
| 247 | Sh3pxd2a | NM 008018 | 1.92741 | 5.16217 | 2.78195 | 2.3669 | 5.45451 | 3.18142 | 2.30711 | 7.95423 | 3.48909 | 2.59193 | 6.03228 | 2.36964 | 0.86281548 |
| 248 | Gm16386 | NR 030709 | 1.19307 | 2.57652 | 3.01428 | 1.48965 | 2.98967 | 2.71596 | 1.22401 | 3.29199 | 3.06786 | 1.89323 | 4.068 | 4.12688 | 0.86117041 |
| 249 | Mllt6 | NM 139311 | 0.735042 | 1.7467 | 1.1979 | 0.6527 | 1.79774 | 1.20146 | 0.648992 | 1.937 | 0.746687 | 0.573013 | 1.74022 | 0.84352 | 0.8520403 |
| 250 | Fgf15 | NM 008003 | 1.06288 | 13.9662 | 8.99268 | 0.677375 | 14.1365 | 8.11848 | 0.559765 | 9.43125 | 6.83936 | 0.357773 | 8.31293 | 5.19388 | 0.84845987 |
| 251 | Fzd4 | NM 008055 | 0.56284 | 1.70361 | 1.40352 | 0.516547 | 1.29787 | 0.813071 | 0.785801 | 1.34871 | 1.01617 | 0.394741 | 1.95267 | 1.19577 | 0.83135929 |

| | | | | | | | | | | | | | | |
|---|---|---|---|---|---|---|---|---|---|---|---|---|---|---|
| 252 | Edn1 | NM 010104 | 0.428122 | 1.71506 | 2.32194 | 0.434988 | 1.25162 | 1.31367 | 0.59032 | 2.53798 | 3.4175 | 0.536083 | 2.14066 | 3.09719 | 0.83007433 |
| 253 | Gse1 | NM 198671 | 1.81401 | 4.64159 | 3.51504 | 1.92311 | 5.59555 | 3.31426 | 1.96535 | 6.19619 | 3.02828 | 1.39511 | 4.93484 | 2.75356 | 0.81740164 |
| 254 | Nrarp | NM 025980 | 2.06781 | 7.06334 | 2.57866 | 1.30333 | 5.96775 | 3.2715 | 1.30223 | 6.02635 | 2.52076 | 1.04405 | 5.40514 | 1.76851 | 0.8153863 |
| 255 | Lzts1 | NM 199364 | 1.56826 | 3.23304 | 3.83774 | 1.69879 | 4.3823 | 3.26689 | 1.0789 | 2.12494 | 2.36385 | 0.459139 | 1.79155 | 2.08761 | 0.7904606 |
| 256 | Bmpr2 | NM 007561 | 2.05186 | 4.86672 | 3.23731 | 1.90771 | 5.03867 | 3.02343 | 2.00556 | 5.36733 | 5.19036 | 1.47838 | 5.42519 | 3.06629 | 0.7899688 |
| 257 | Tbx3 | NM 011535 | 1.5167 | 3.83349 | 2.39045 | 0.655665 | 4.48431 | 1.0497 | 0.493388 | 3.77273 | 2.48705 | 1.09363 | 5.43874 | 2.17601 | 0.74224122 |
| 258 | Pdgfb | NM 011057 | 0.418682 | 3.46743 | 2.0801 | 0.529958 | 2.99701 | 1.43711 | 0.210024 | 2.39491 | 1.51745 | 0.282163 | 2.69393 | 1.76146 | 0.66523418 |
| 259 | Cbln4 | NM 175631 | 0.47797 | 1.72927 | 2.16482 | 0.42985 | 1.54504 | 2.31995 | 0.140651 | 1.03924 | 1.61484 | 0.317851 | 1.16015 | 1.348 | 0.65336262 |
| 260 | T | NM 009309 | 0.383501 | 2.10844 | 3.93727 | 0.496742 | 1.95387 | 3.65809 | 0.145134 | 1.4556 | 2.64447 | 0.378694 | 1.89654 | 4.14447 | 0.62718331 |
| 261 | Mafb | NM 010658 | 0.414879 | 2.79711 | 1.45158 | 0.74214 | 2.21235 | 1.13129 | 0.282065 | 1.58468 | 1.07413 | 0.18156 | 1.94538 | 0.744864 | 0.59530186 |
| 262 | Gata4 | NM 008092 | 0.745822 | 2.24336 | 0.478218 | 0.879517 | 2.82786 | 0.773617 | 0.188194 | 1.57141 | 0.699541 | 0.762483 | 1.95403 | 0.485598 | 0.570268 |
